# Dynamic allocation of carbon storage and nutrient-dependent exudation in a revised genome-scale model of *Prochlorococcus*

**DOI:** 10.1101/2020.07.20.211680

**Authors:** Shany Ofaim, Snorre Sulheim, Eivind Almaas, Daniel Sher, Daniel Segrè

**Affiliations:** Bioinformatics Program and Biological Design Center, Boston University, Boston, MA, USA; Department of Marine Biology, University of Haifa, Haifa, Israel; Department of Biotechnology and Food Science, NTNU-Norwegian University of Science and Technology, Trondheim, Norway; Department of Biotechnology and Nanomedicine, SINTEF Industry, Trondheim, Norway; K.G. Jebsen Center for Genetic Epidemiology, NTNU-Norwegian University of Science and Technology, Trondheim, Norway; Department of Biomedical Engineering, Boston University, Boston, MA; Department of Physics and Department of Biology, Boston University, MA

## Abstract

Microbial life in the oceans impacts the entire marine ecosystem, global biogeochemistry and climate. The marine cyanobacterium *Prochlorococcus*, an abundant component of this ecosystem, releases a significant fraction of the carbon fixed through photosynthesis, but the amount, timing and molecular composition of released carbon are still poorly understood. These depend on several factors, including nutrient availability, light intensity and glycogen storage. Here we combine multiple computational approaches to provide insight into carbon storage and exudation in *Prochlorococcus*. First, with the aid of a new algorithm for recursive filling of metabolic gaps (ReFill), and through substantial manual curation, we extended an existing genome-scale metabolic model of *Prochlorococcus* MED4. In this revised model (*i*SO595), we decoupled glycogen biosynthesis/degradation from growth, thus enabling dynamic allocation of carbon storage. In contrast to standard implementations of flux balance modeling, we made use of forced influx of carbon and light into the cell, to recapitulate overflow metabolism due to the decoupling of photosynthesis and carbon fixation from growth during nutrient limitation. By using random sampling in the ensuing flux space, we found that storage of glycogen or exudation of organic acids are favored when the growth is nitrogen limited, while exudation of amino acids becomes more likely when phosphate is the limiting resource. We next used COMETS to simulate day-night cycles and found that the model displays dynamic glycogen allocation and exudation of organic acids. The switch from photosynthesis and glycogen storage to glycogen depletion is associated with a redistribution of fluxes from the Entner-Doudoroff to the Pentose Phosphate pathway. Finally, we show that specific gene knockouts in *i*SO595 exhibit dynamic anomalies compatible with experimental observations, further demonstrating the value of this model as a tool to probe the metabolic dynamic of *Prochlorococcus*.

## Introduction

Marine phytoplankton perform about one-half of the photosynthesis on Earth (Field et al., 1998). *Prochlorococcus* is one of the most abundant phytoplankton clades in the world’s oceans and is estimated to produce about 4 Gt of organic carbon annually (Flombaum et al., 2013a). As such, these clades play a key role in a variety of ecosystems (reviewed by (Partensky and Garczarek, 2010; Biller et al., 2015)). Recent evolutionary studies suggested several evolved metabolic innovations contributing to high picocyanobacterial abundance in the harsh oligotrophic ocean waters, (Field et al., 1998) usually limited by several nutrients such as nitrogen, phosphorus, and iron. These innovations include a proteome that contains less nitrogen rich amino acids (Gilbert and Fagan, 2011), membranes that contain glyco- and sulfolipids rather than phospholipids (Van Mooy et al., 2006) and streamlining of the genome associated with outsourcing of important cellular functions to co-occurring organisms (Holtzendorff et al., 2008; Partensky and Garczarek, 2010; Morris et al., 2012; Ma et al., 2018a; Braakman, 2019).

Another innovation employed by these organisms is an increased metabolic rate that in turn manifest in the exudation of organic compounds (Braakman et al., 2017; Braakman, 2019), resulting in typically 2–50% of the carbon fixed by photosynthesis leaving the cell (Kujawinski, 2011; Thornton, 2014). Consequently, dissolved organic matter is an omnipresent component of natural waters. However, it is currently impossible to provide a universal chemical description of the dissolved organic matter (Kujawinski, 2011; Arrieta et al., 2015; Moran et al., 2016), partly because the exuded organic compounds differ between strains and environmental conditions (e.g. (Becker et al., 2014; Ma et al., 2018b)). Nevertheless, in general, phytoplankton exudate includes a small proportion of low-molecular weight compounds, such as organic acids, carbohydrates, and amino acids (Bertilsson and Berglund, 2014), as well as a larger proportion of complex, high-molecular weight compounds (Kujawinski, 2011). Another strategy employed by these bacteria to manage their carbon budget is the internal storage of carbon in polymeric form, specifically, glycogen (Zinser et al., 2009; Reimers et al., 2017; Luan et al., 2019; Szul et al., 2019). Glycogen accumulates in the bacterial cell during the light hours and was recently suggested to have two primary roles; as energy storage in preparation for darkness and as a regulation strategy to manage high-light photosynthesis byproducts (Welkie et al., 2019). The allocation of glycogen is suggested to be tightly associated with the overflow metabolism hypothesis and also known to be widely affected by nutrient limitations (Damrow et al., 2016; Cano et al., 2018; Forchhammer and Schwarz, 2019; Szul et al., 2019). Importantly, the carbon fixed and released by phytoplankton is then used by heterotrophic organisms as a source of energy, whereas the heterotrophic bacteria may recycle nutrient elements and support the growth of phytoplankton in other ways, as suggested by the Black Queen Hypothesis (Amin et al., 2012; Morris et al., 2012; Moran et al., 2016; Cirri and Pohnert, 2019; Moran and Durham, 2019). Thus, carbon fixation, storage and release are tightly intertwined with microbial interactions and microbial ecosystem dynamics.

Quantitative models at various scales have provided critical insights into how ocean microbial ecosystems function, and how they are related to broader biogeochemical cycles (Braakman et al., 2017; Coles et al., 2017; Foster et al., 2018; Moradi et al., 2018). Most of these models represent organisms in terms of simplified stoichiometric reactions converting elements into biomass, thus making it possible to incorporate biological processes into dynamic-coupled Earth System models (Follows et al., 2007; Reid, 2011). The exponential increase in genomic information on marine organisms provides an opportunity to seek methods to link such detailed genome-scale information to biochemical flows (Coles et al., 2017). In recent years, genome-scale metabolic models (GEMs), combined with linear programming, have made it possible to produce testable predictions of metabolic phenotypes of individual organisms or microbial communities (Gu et al., 2019). This computational framework is based on the identification of individual enzymes and transporters in an organism’s genome, and on simplifying assumptions that bypass the need for kinetic parameters (Maarleveld et al., 2013; O’Brien et al., 2015; Casey et al., 2016; Kim et al., 2016; Reimers et al., 2017). While genome-scale modeling has proven to be a powerful approach in cyanobacterial model organisms such as *Synechocystis* sp. PCC 6803, *Synechococcus* elongatus PCC 7942 and *Prochlorococcus marinus* MED4 (Knoop et al., 2013; Broddrick et al., 2016; Casey et al., 2016; Yoshikawa et al., 2017), the exudation of organic compounds in phototrophic organisms has not been studied in detail through Flux Balance Analysis (FBA) or similar methods (Varma and Palsson, 1994; Orth et al., 2010). On the other hand, several examples exist of FBA-based predictions of exudation-mediated interactions between different species, including using the COMETS platform (Harcombe et al., 2014). In fact, FBA calculations also suggest that “costless” secretions (i.e. secretions that do not induce a fitness cost) might be quite common, and can support the growth of co-occurring organisms (Pacheco et al., 2018).

Experimental evidence and theoretical considerations indicate that *Prochlorococcus* exudes different metabolites in a way that strongly depends on environmental conditions (Dubinsky and Berman-Frank, 2001; Szul et al., 2019) as well as on the strain’s genetic makeup (Becker et al., 2014; Roth-rosenberg et al., 2019). While GEMs can be used to predict these fluxes, they require modifications to deal with processes not usually considered in FBA, including: (a) the special nature of photon fluxes (which, unlike molecular fluxes, cannot easily be “shut off” at short time scales, (Dubinsky and Berman-Frank, 2001)); (b) the buffering role of intracellular storage molecules such as glycogen. The primary focus of this study is to obtain better knowledge of the potential metabolic effect of a combination of key nutrients (carbon, nitrogen, phosphorous and light) and carbon fixation rate on the allocation (including storage and exudation) of carbon in *Prochlorococcus* using a revised genome-scale metabolic model (Figure 1). We start by describing model revisions and updates to capture the current, most complete metabolic knowledge available for *Prochlorococcus*. Next, we use a variety of FBA approaches to uncover the potential relationships between a set of key nutrients, carbon storage and exudates in static and dynamic (time dependent) settings. The implementation and use of these approaches improve our understanding of the intricate metabolic workings of *Prochloroccoccus* and provide insights on its storage and exudation trends under different environmental conditions.

**Figure 1:**
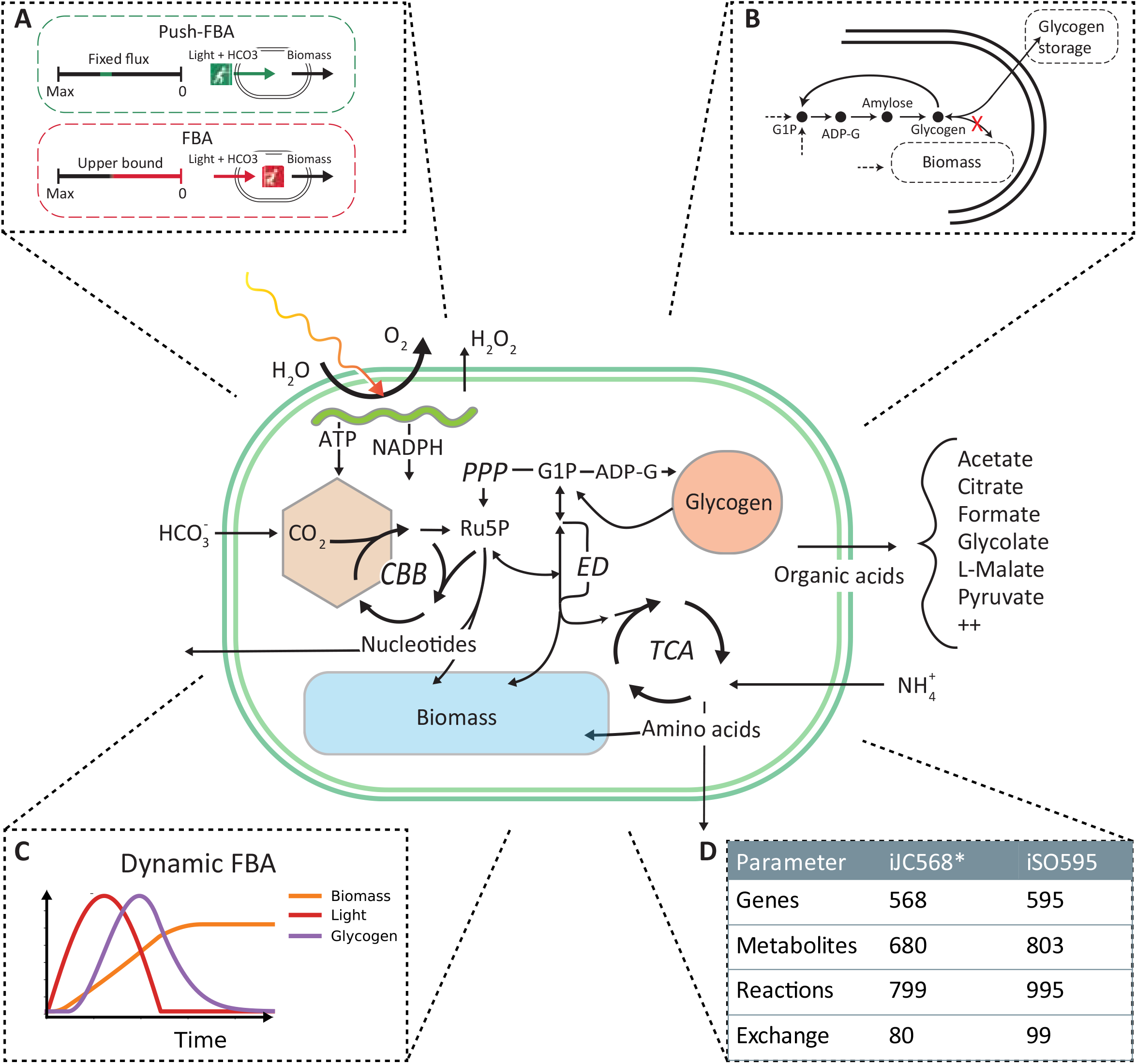
iSO595 is an updated reconstruction of *P. marinus* MED4, featuring a complete Entner-Doudoroff pathway, rewired glycogen metabolism and increased coverage of the genome. The central panel is a simplified illustration covering the most relevant metabolic features. **A)** To simulate the natural environmental constraints experience by Prochlorococcus we use “Push-FBA”: Light and bicarbonate uptake is given a fixed flux independent of growth rate. This contrasts standard FBA where light and bicarbonate would be “pulled” in by the demand needed to support maximal growth rate up to a given bound. **B)** We rewired the glycogen metabolism in iSO595 to study the dynamic allocation the dynamic allocation of glycogen. **C)** Additionally, we implemented dynamic light conditions and light absorption in COMETS to simulate the growth of *P. marinus* during the diel cycle. **D)** iSO595 has increased coverage both in terms of genes, reactions and metabolites compared to its ancestor iJC568.

## Methods

### Model update and curation

The *i*JC568 genome-scale reconstruction of *Prochlorococcus marinus subsp. pastoris str*. CMP1986 (referred to throughout the manuscript as MED4) as described by Casey et al. (2016), was used as the starting point for model enhancement. The update process started with an in-depth study of the reconstructed network and available knowledge not previously incorporated into the model of the organism. During this process, we ended up implementing the following specific steps of curation and update: (i) A key modification to the model was the decoupling between the glycogen storage flux and the biomass production. In standard stoichiometric reconstructions for FBA modeling (Thiele et al., 2011; Nogales et al., 2012; Feist et al., 2014; Broddrick et al., 2016; Monk et al., 2017; Kavvas et al., 2018), glycogen is listed as one of the biomass components, thus accounting for the carbon flux into storage. However, given the fixed stoichiometry of biomass composition, this classical implementation cannot account for the time-dependent storage and re-utilization of glycogen observed in picocyanobacteria. We thus removed the glycogen from the biomass function and streamlined the existing glycogen granule representation to a direct link between ADP-Glucose to the production of glycogen (Figure 1B). (ii) In addition to targeted refinement of selected reactions, we used the KEGG database (Kanehisa and Goto, 2000) to perform an extensive search of previously known but missing metabolic reaction annotations. Indeed, we found 354 reactions that could be potentially added to the existing network. To incorporate this knowledge, we developed a semi-automated algorithm (ReFill, described below), (iii) We coupled the implementation of the algorithm with several steps of manual curation. These included the addition of transports, such as that of hydrogen peroxide and ethanol, known to diffuse across the cell membrane (Seaver and Imlay, 2001; Noreña-Caro and Benton, 2018), and the addition of the complete Entner-Doudoroff pathway, that has recently been discovered in cyanobacteria (Chen et al., 2016). Additionally, we performed a BLAST search (Supplemental material 1) (Altschul et al., 1990) from which we identified 6PG-dehydratase (EC: 4.2.1.12) encoded by PMM0774, thereby completing this pathway in the model reconstruction. (iv) The revised model was checked for redox and elemental balance. Since the biomass function was based on experimental data (Casey et al., 2016), it was not updated. In line with best practices, a *memote* quality assessment (Supplemental material 2) (Lieven et al., 2020), as well as model files and a detailed changelog, are provided at https://github.com/segrelab/Prochlorococcus_Model. All reactions added to iJC658 to form iSO595 are found in Table S1 and modified reactions are found in Table S2.

### ReFill algorithm

Following an extensive search of literature and the KEGG (Kanehisa and Goto, 2000), TransportDB (Elbourne et al., 2017) and Metabolights (Haug et al., 2013) databases, we found a large number of new or previously known but missing reaction, transporter and metabolite annotations. Adding large amounts of data to an existing network might create new gaps and may give rise to new blocked reactions and orphan metabolites. These, in turn, may affect optimization simulations. To add this knowledge to the network in a controlled approach, we developed the semi-automated recursive algorithm ReFill (Recursive Filler (of metabolic gaps)). The algorithm is based on the principle of using diverse information, such as enzyme and reaction annotations, and experimental data (such as metabolomics), to selectively increase the metabolic knowledge of an organism’s existing curated genome-scale metabolic network. ReFill makes use of a repository of reactions, in this case KEGG reaction annotations for MED4 absent from the model, to construct all potential chains of reactions connecting two metabolites in the existing network. It systematically tests the potential of adding each new reaction and suggests adding it only if it can be a part of a chain in which all the metabolites are part of a path in the network (Figure 2). This prevents the creation of new orphan metabolites and potential blocked reactions that may lead to infeasible FBA solutions. The algorithm starts by selecting a reaction from the repository. It then inspects each metabolite in the reaction for presence in the existing network. In case a metabolite is not present, the set of available reactions is scanned for other reactions using this metabolite as a substrate or product. If such a reaction is found, it is added to the chain of potential reactions. The algorithm then iteratively expands the chain until either the repository is exhausted or all the metabolites in the most recent reaction added are present in the network. After all the possible chains of new reactions are expanded, the algorithm examines the connectivity of all the metabolites in each chain (See example in Figure 2B). Following the manual addition of transporters found through TransportDB (Elbourne et al., 2017) and Metabolights (Haug et al., 2013) (Study MTBLS567), using the ReFill algorithm, we updated reactions that belong to several different pathways, including metabolism of cofactors and vitamins, carbohydrate metabolism, amino acid metabolism and nucleotide metabolism. A complete list of added reactions can be found in Table S1. ReFill was coded in python 3.7 and generates MATLAB-compatible files formatted to be used with the COBRA MATLAB toolbox, including a list of suggested reactions to add and their gene-reaction rules. Other outputs include the added reaction chains and possible metabolic circuits that can be formed by these additions. All code (including ReFill) and model files are available on https://github.com/segrelab/Prochlorococcus_Model.

**Figure 2:**
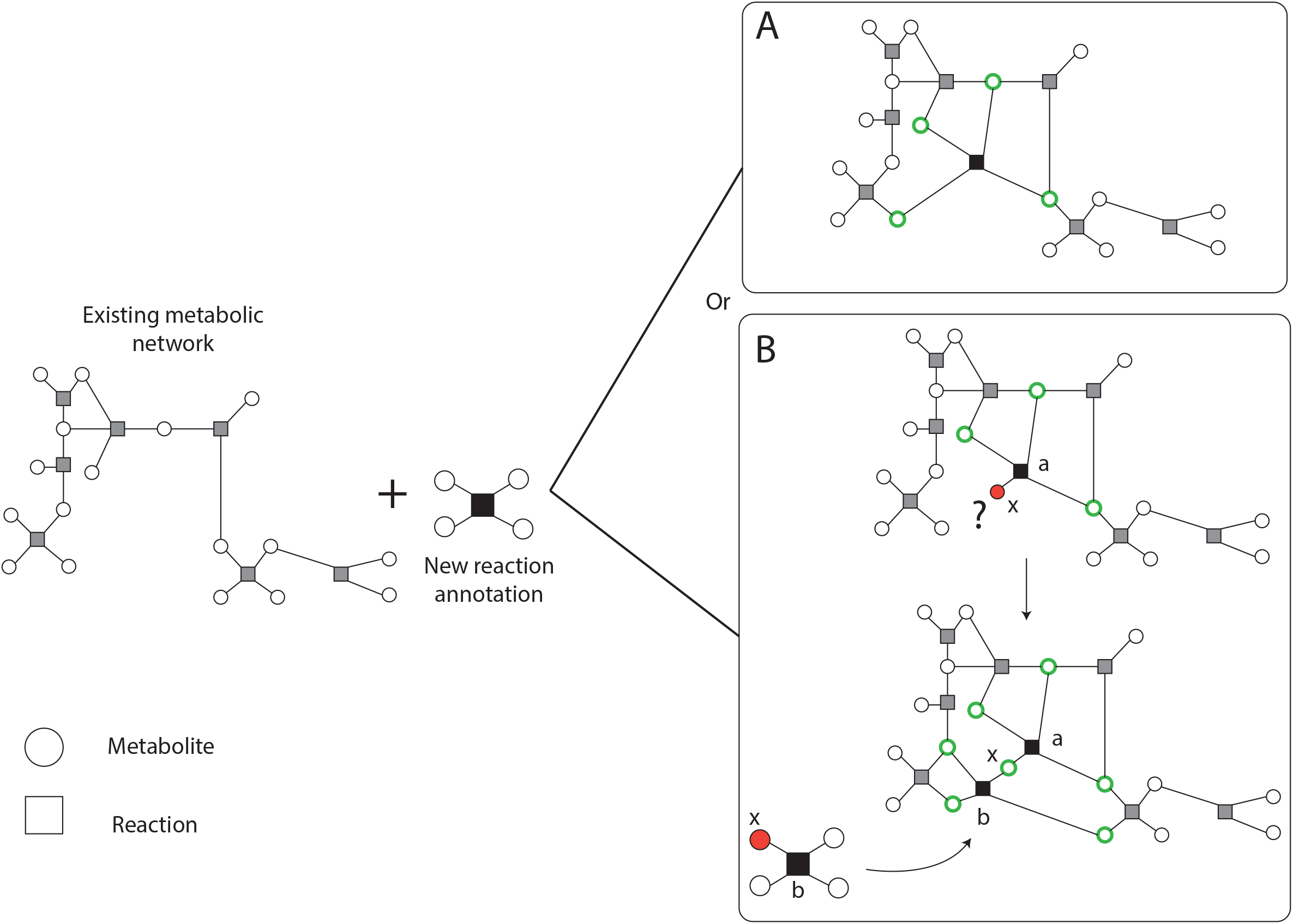
Schematic description of the ReFill algorithm. New reaction annotations are added only if all the metabolites are connected to the existing network. A connected metabolite is denoted in a green circle. An orphan (unconnected metabolite is denoted in a red circle) (A) A simple case in which all the metabolites comply with the selection rule. (B) In case one or more metabolites are not connected to the existing network, other new reactions may be added to complete the missing connections. For example, consider reaction a to be composed of two substrates and two products, one of which, x (a metabolite that did not exist in the initial network), is not connected to the existing network and is currently an orphan. After the expansion step, the algorithm identified one reaction, b, in which x is used as a substrate. All other metabolites in b exist in the network thus creating a path from reaction a through reaction b to the network.

### Parameter sampling

To study the effects of combinations of key nutrients on glycogen production and exudation in the *i*SO595 model we focused on four parameters representing the uptake fluxes of light, bicarbonate, ammonium, and phosphate. Light and inorganic carbon (bicarbonate) are the substrates for photosynthesis, whereas nitrogen and phosphorus limit the growth of *Prochlorococcus* in large regions of the world ocean (Davey et al., 2008; Moore et al., 2013; Saito et al., 2014), and nutrient limitation is likely to influence the exudation of fixed carbon (Dubinsky and Berman-Frank, 2001). We sampled 10,000 different environmental conditions by drawing random values from uniform distributions of these four parameters. The range of each parameter was based on physiologically relevant ranges we extracted from the literature and on the requirement that each range covers important phase transitions, such as nutrient and light limitations (Table S3). Light flux was converted from micromole quanta m^−2^ sec^−1^ to mmol gDW^−1^h^−1^ similarly to (Nogales et al., 2012) using 8% photosynthesis efficiency rate (Zinser et al., 2009). All parameter uptake fluxes are described in FBA compatible units (mmol gDW^−1^h^−1^), corresponding values in biogeochemistry relevant units are illustrated in Table S3. As MED4 is a photoautotroph, it is exposed to a constant stream of light during daylight hours. The bacterium is then forced, or ‘pushed’, to fix carbon even when there is not enough of other elements, such as nitrogen or phosphate, to combine the fixed carbon into biomass. To capture this phenomenon *in silico* we developed a ‘push’-FBA framework where we fixed both upper and lower uptake rates of light and bicarbonate (Figure 1A). For the other sampled nutrients, ammonium and phosphate, we defined standard FBA bounds where the maximal uptake rate was set to the sampled value and the lower bound was set to zero. Note that, we only considered uptake of sulfur in the form of sulfate (not hydrogen sulfide), and no upper limit was set for the uptake of sulfate because of its abundance in seawater. The maximum rate of RuBisCO (R00024) was fixed to 4.7 mmol gDW^−1^ h^−1^, as previously reported (Casey et al., 2016). Before sampling we blocked a set of artificial exchange reactions that were added in the previous version of the model, most likely to allow export of dead-end metabolites that would otherwise limit flux feasibility (Table S4). Subsequently, we removed all unconditionally blocked reactions in the model to speed up computations. For each random sample, we first tested the model for feasibility using FBA (Varma and Palsson, 1994). If the solver returned a solution that was feasible and optimal, we further calculated optimal fluxes with parsimonious FBA (Lewis et al., 2010), and determined the range of possible fluxes at optimum with Flux Variability Analysis (FVA) (Gudmundsson and Thiele, 2010). Exchange fluxes from FBA, parsimonious FBA and FVA were recorded and used in subsequent analyses. All environmental sampling and calculations were performed using CobraPy (Ebrahim et al., 2013) and GUROBI 8.1.1 (Gurobi Optimization, Inc., Houston, TX, USA). The python script used to run the environmental sampling is available in the GitHub repository (https://github.com/segrelab/Prochlorococcus_Model).

### Statistical analysis of the sampled spaces

We sampled 10,000 different environmental conditions based on the flux ranges described above, and analyzed the results of FBA optimization, with the goal of characterizing the distribution of specific import/export fluxes and the correlations between different fluxes. To that end, we calculated Pearson correlations between reaction fluxes in the sampling data using the python (version 3.7) Pandas package version 1.0.3 (McKinney, 2010). While negative values are normally used to define uptake in FBA, we converted them to positive values for the uptake of light, bicarbonate, phosphate, ammonium, and sulfate when calculating correlations to ease interpretation of the results. We also performed hierarchical clustering using the Nearest Point Algorithm in SciPy (Virtanen et al., 2020) to sort the order of the compounds in the correlation matrix. We performed dimensionality reduction on normalized reaction fluxes using the T-distributed Stochastic Neighbor Embedding (t-SNE) method (Maaten and Hinton, 2008) in Scikit-learn (Pedregosa et al., 2011) with perplexity of 50 and 3000 iterations. The reaction fluxes were normalized to [-1,1] by dividing by the maximum absolute flux of each reaction. Finally, the t-SNE transformed data was clustered using HDBSCAN (McInnes et al., 2017) with a minimum cluster size of 200. Transport of inorganic ions, water, and protons were not considered when calculating correlations, dimensionality reduction or clustering. We also discarded transport reactions with no absolute flux value above 10^−3^ mmol gDW^−1^ h^−1^ in any of the environmental samples.

### Dynamic modelling of light absorption during the diel cycle in COMETS

Cyanobacteria follow a diel cycle. To capture this dynamic behavior, we extended the Computation Of Microbial Ecosystems in Time and Space (COMETS) platform (Harcombe et al., 2014), and developed a module for diurnal-cycle simulations allowing oscillations of light intensity and light absorption. Attenuation of light through each grid cell was modelled using the Beer-Lambert law, as described previously (Yang, 2011; Gomez et al., 2014):

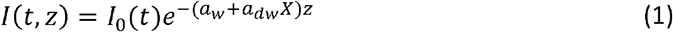

Here, *I*(*t, z*) is the light irradiance given in mmol photons m^−2^ s^−1^, *t* is the time, *z* is the depth (from the top of the grid cell), *a_dw_* is the cell- and wavelength-specific absorption coefficient given in m^2^ gDW^−1^, *a_w_* the absorption coefficient of pure water given in m^−1^, *X* the biomass concentration in gDW m^−3^, and *I*_0_(*t*) time time-dependent incident light irradiance at the top of the grid cell. In the current version, we simplified the process by assuming that the light irradiance is either monochromatic or a sum of the total light bandwidth, and the absorption coefficient should match the wavelength(s) of the light source. The total light attenuation (Δ*I*) through a grid cell of thickness Δ*z* is then

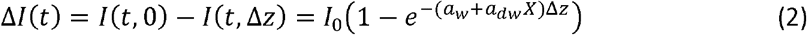

The light absorbed by the cells is a fraction of the total light attenuation, i.e.

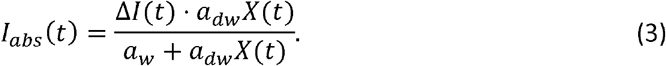

The total number of photons absorbed per dry cell weight (Φ(*t*)) in mmol photons gDW^−1^ s^−1^ by the cells within a grid cell of thickness Δ*z*, volume *V* and surface area *A* is then

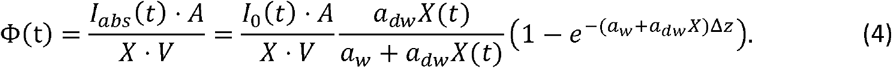

For all COMETS simulations presented here we have used monochromatic light at 680 nm with a calculated biomass-specific absorption coefficient *a_dw_* as previously described (Morel and Bricaud, 1981; Bricaud et al., 2004). Briefly, the biomass-specific absorption is the weighted sum of the absorption coefficients of the light-absorbing pigments divinyl-chlorophyll A and B, since none of the other pigments in *Prochlorococcus* absorb light at 680 nm. Additionally, to account for the discrete distribution of chlorophyll into separate cells, the absorption coefficient is scaled by the packaging factor. All coefficients used to calculate light attenuation and absorption are provided in Table 1.

**Table 1:** Coefficients and values used to calculate light absorption in COMETS.

| Symbol | Description | Value | Unit | Reference |
| --- | --- | --- | --- | --- |
| $\lambda$ | Wavelength | 680 | nm | |
| $a_w$ | Absorption coefficient of water at 680 nm | 0.465 | m<sup>-1</sup> | Pope and Fry, 1997 |
| $a_{dw}$ | Biomass-specific absorption coefficient | 0.285 | m<sup>2</sup> gDW<sup>-1</sup> | |
| **a<sub>dvchl-A</sub>** | Absorption coefficient of divinyl-chlorophyll A at 680 nm | 0.0184 | m<sup>2</sup> (mg dvchl-A)<sup>-1</sup> | Bricaud et al., 2004 |
| **a<sub>dvchl-B</sub>** | Absorption coefficient of divinyl-chlorophyll B at 680 nm | 0.0018 | m<sup>2</sup> (mg dvchl-B)<sup>-1</sup> | Bricaud et al., 2004 |
| **c<sub>dvchl-A</sub>** | Amount of divinyl-chlorophyll A in | 0.0163 | g dvchl-A gDW<sup>-1</sup> | Casey et al., 2016 |
| **c<sub>dvchl-B</sub>** | Amount of divinyl-chlorophyll B | 0.0013 | g dvchl-B gDW<sup>-1</sup> | Casey et al., 2016 |
| **d** | Average diameter of MED4 | 0.6 | μm |  |
| **n'** | Imaginary part of the refractive index at 675 nm | 0.01377 |  | Stramski et al., 2001 |
| **Q\*** | Packaging effect at 680 nm | 0.945 |  |  |

The changing light conditions throughout a diel cycle was modelled as

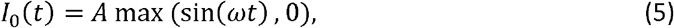

where the angular frequency is 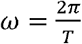.

Following the development of the diel cycle simulation capability in COMETS we set out to dynamically simulate the growth of MED4. Since the nutrient uptake follows Michaelis–Menten kinetics, we estimated the kinetic parameters *V_max_* and *K_m_* using a heuristic approach from experimental data (Grossowicz et al., 2017), first by finding the range of possible parameter combinations corresponding to the gross growth rate of 0.5 d^−1^ (Figure S1A), and secondly by comparing predicted growth and ammonium depletion with the experimental time-series cultivation data (Figure S1B). The estimated parameters were used in the remaining dynamic FBA simulations in COMETS. Finally, to simulate the dynamic storage and consumption of glycogen we applied a multiple objective approach consisting of the following four steps: (1) Maximization of the flux through the non-growth associated maintenance reaction. Note that, this reaction has an upper bound of 1 mmol gDW^−1^ h^−1^ (Casey et al., 2016) In contrast to standard practice, where one uses a lower bound for the non-growth associated maintenance reaction, this method provides a more realistic scenario where the organism continues to consume resources trying to keep up cellular maintenance even at zero growth; (2) Maximization of growth; (3) Maximization of glycogen production (storage); and (4) Parsimonious objective which minimizes the sum of absolute fluxes. To simulate nitrogen-abundant and nitrogen-poor growth conditions, we used the PRO99 medium with standard (800 μMol) and reduced (100 μMol) ammonium concentration, as previously described (Grossowicz et al., 2017). Light availability was modelled as described in Equation 5, with an amplitude of 40 μmol Q m^−2^ s^−1^ and a period of 24 hours. All parameter values used in the COMETS simulations are given in Table S5. All Dynamic growth simulations were performed using COMETS v.2.7.4 with the Gurobi 8.1.1 solver, invoked using the associated MATLAB toolbox (https://github.com/segrelab/comets-toolbox).

### Simulating growth of knockout mutants

Simulations of the knockout mutants where performed by constraining the flux to zero for the reactions catalyzed by the enzymes encoded by *glgC* (PMM0769) and *gnd* (PMM0770), respectively. For *glgC*, the reaction is glucose-1-phosphate adenylyltransferase (R00948) and for *gnd* the two reactions are NADP^+^ and NAD^+^ associated 6-phosphogluconate dehydrogenases (R01528 and R10221). We then used dynamic FBA in COMETS with PRO99 medium (Moore et al., 2007) with limited ammonium and diel light conditions to simulate growth over 7 days. We included an arbitrarily derived death rate of 0.1 d^−1^ (Grossowicz et al., 2017) in these simulations. The growth curves where qualitatively compared with experimental data from Shinde and co-workers (Shinde et al., 2020). The model parameters are given in COMETS parameter and model files in https://github.com/shashany/iSO595/tree/master/COMETS_files.

## Results and Discussion

### Model curation and update

*Prochlorococcus* fixes carbon through photosynthesis during daytime. Fixed carbon that is neither used for cell growth nor stored in the form of glycogen is exuded. Here, we set out to study dynamic changes in the carbon allocation and storage mechanisms in MED4 using a genome-scale metabolic modelling approach. To that end, we first re-curated and updated the available *i*JC586 model (Casey et al., 2016), as described in detail in the Methods section. The update involved the development of a new semi-automatic algorithm (ReFill), which can be broadly applied to other reconstructions (Methods). Concurrently, we introduced a revised mechanism for carbon storage, effectively treating glycogen as an independent component of biomass. This configuration makes it possible for glycogen to be accumulated and depleted at variable rates (Figure 1), aligning with the overflow metabolism hypothesis (Szul et al., 2019; de Groot et al., 2020). Other key modifications induced by the ReFill algorithm and subsequent manual curation (see Methods) include the completion of the Entner-Doudoroff (ED) pathway, recently discovered in cyanobacteria (Chen et al., 2016) and proposed as the primary *Prochlorococcus* glucose metabolism pathway under mixotrophic conditions (Biller et al., 2018), but see (Muñoz-Marín et al., 2020). Additional revisions focused on the exudation of fixed carbon products from the cell and included various transports such as pyruvate, fumarate, citrate, ethanol, various nucleotides and hydrogen peroxide as well as metabolites found in both the endo- and exo-metabolome of *Prochlorococcus* (Metabolights study MTBLS567). The end product of our revision, reconstruction iSO595, has 595 genes, 802 metabolites and 994 reactions, i.e. 27 genes, 123 metabolites and 196 reactions more than the previous version, *i*JC568 (Figure 1D).

### Carbon fixation and storage is affected by nutrient uptake rate

*Prochlorococcus* thrive in oligotrophic environments (Johnson et al., 2006), where, in surface waters, its growth and carbon fixation rates are usually limited by the abundance of nitrogen, phosphate or iron (Krumhardt et al., 2013; Szul et al., 2019). Deeper in the water column *Prochlorococcus* growth becomes limited by light (Vaulot et al., 1995). We set out to explore the combined effect of different levels of light and nutrients on carbon fixation, storage and exudation. Similarly to Phenotypic Phase Plane analysis (Edwards et al., 2002), we sought a global perspective of metabolism in this multi-parameter spaces while explicitly taking into account the fact that the inflow of light and bicarbonate may not be easily controllable by the cell, and that *Prochlorococcus* may need to deal with excess amounts of fixed carbon. Thus, in contrast to normal FBA where the uptake of metabolites is constrained by an upper bound, we introduced a ‘push-FBA’ approach (Figure 1A), in which the influx of bicarbonate and light have a fixed imposed value (See Methods and Table S3 for specific values used). This approach attempts to mimic implications of photosynthesis, in which light is the driving force. Once photons are absorbed by the chlorophyll, the energy must be used to produce ATP and reducing power, otherwise it is dissipated in ways that may cause cell damage (Long et al., 1994). This subtle difference in applied constraints has major effects on model predictions. While flux rearrangement is usually viewed as a consequence of environmental nutrient limitations, the results of this analysis shows that a substantial rewiring of fluxes is caused by this imposed excess of fixed carbon as well.

To understand how different combinations of environmental parameters (availability of nitrogen, phosphate, light and bicarbonate) affect the way *Prochlorococcus* can manage its carbon budget, we implemented FBA under 10,000 randomly sampled growth environments. Overall, this sampling analysis demonstrated that the exudation of organic acids, amino-acids and nucleobases/nucleosides, as well as the extent of glycogen storage, are strongly modulated by environmental factors (Figure 3). To observe the full range of possible optimal solutions per sample, we implemented and compared different flux balance analysis methods, including flux variability analysis (FVA) and parsimonious FBA (pFBA). These two methods provide complementary insight: FVA estimates the range of possible values for the flux of each reaction at the optimum, providing insight into the structure of the phenotypic space at maximal growth rate. In contrast, pFBA, by minimizing the sum of fluxes at optimality, generates flux predictions less likely to involve unrealistic loops, and thus potentially provides predictions closer to experimental values (Lewis et al., 2010). Together, these two FBA methods help analyze the solutions of our high-dimensionality dataset.

**Figure 3:**
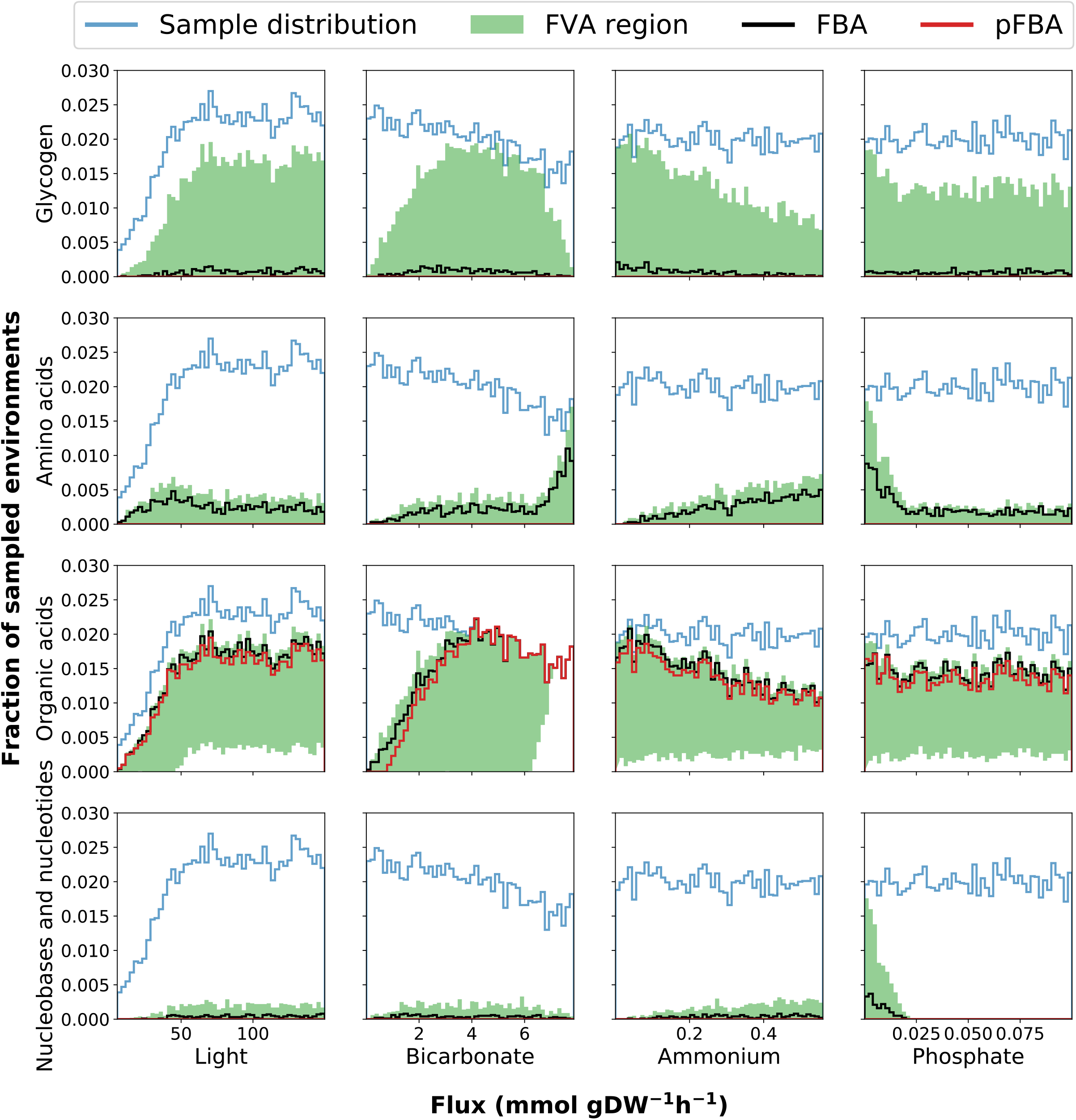
Histograms of environmental sampling results provide insight into how the fixed uptake rates of light and bicarbonate and the upper bounds on ammonium and phosphate affects exudation of organic acids, nucleobases and nucleotides, amino acids as well as glycogen storage. The y-axis represents the fraction of all sampled environments yielding feasible models. Although we initially sample each parameter uniformly the final sample distribution is not uniform because some combination of parameters represents infeasible phenotypes (no solution can satisfy the constraints). The final sample distribution for each parameter is therefore shown in the figure panels as a blue line. The black and red line represents the histograms of samples where exudation is predicted by FBA and pFBA, respectively. The shaded green region represents the span between histograms of samples as predicted by FVA: the lower and upper bound represents the number of samples where exudation is predicted using the minimum and maximum value of FVA, respectively. This FVA region covers the range of possible phenotypes. The lower bound of the FVA region display the number of samples where a certain outcome is obligatory to maximize growth, while the upper bound of the FVA region display the number of samples where the outcome is possibly without reducing growth.

Our predictions simulate the metabolic effects and variability in glycogen production modulated by environmental constraints (Figure 3). Glycogen production was observed only above light levels of 50 mmol gDW^−1^ h^−1^ (corresponding to 7.5 micromole quanta m^−2^ sec^−1^), and decreased as ammonium and phosphate concentrations increase, in agreement with previous studies (Szul et al., 2019). Interestingly, FVA consistently predicted the glycogen production range minimal value to be zero across all samples. This implies that glycogen storage is possible, but not necessary to achieve optimal growth in the feasible solution space. This was also the case in the more stringent pFBA analysis, indicating that while metabolism may be a strong modulator of glycogen metabolism, more types of regulation, not accounted for in FBA, are involved. One example of such regulation may be allosteric regulation of ADP-glucose pyrophosphorylase by 3-phosphoglycerate (Iglesias et al., 1991), possibly in combination with redox regulation (Díaz-Troya et al., 2014). Specific regulation aimed at tuning up glycogen storage may also occur at the transcriptional level, e.g. by multiple transcription factors previously suggested to be involved in the regulation of glycogen metabolism in fluctuating environments (Luan et al., 2019).

The range of possible rates of glycogen production (through FVA) displays a bell-shaped bicarbonate dependent distribution, indicating low storage of glycogen (zero flux) under both low and high uptake rates of bicarbonate. When bicarbonate uptake rates are low, all available carbon is diverted into growth. The reduced glycogen storage at high bicarbonate uptake, when RuBisCO is saturated, seems to be caused by the increased ATP demand associated with the conversion of bicarbonate to exudation-products, since the onset and rate of change of this trade-off is modulated by the ATP availability, as demonstrated by phenotypic phase planes analysis (Figure S2). This agrees with recent work suggesting that *Prochlorococcus* use available ATP to drive pathways to saturation by shifting reaction directions towards favoring dephosphorylation of ATP to ADP, disrupting the cellular ATP/ADP ratio and increasing the metabolic rate of the cell by pushing forward ATP consuming reactions, until it is restored. Together with organic carbon exudation this strategy allows for growth in lower nutrient concentrations (Braakman, 2019).

We next sought to explore the effect of combinations of key nutrients on storage and exudation patterns in our sampling spaces. To explore the strongest trends, we chose to employ a high stringency approach and use only our set of pFBA results. Due to the nature of pFBA, any exudation observed in this analysis could not be easily removed without imposing a cost on growth. To that end, we visualized the data using t-SNE clustering (Figure 4A) on our pFBA set of results. We observed 6 typical phenotypes (clusters) rising out of the sampling spaces (Figure 4). These 6 phenotypes are characterized by subtle differences in combinations of environmental parameters, yielding significantly different exudation patterns. Generally, we observed the highest biomass value in phenotype 5, and the lowest in phenotype 4. All key nutrient uptake rates were highly variable (ranging from 33 to 44% variability). Phenotype 1 is characterized by high light, bicarbonate, a maximum RuBisCO flux (indicating maximal photosynthesis rate) but low nitrogen uptake. Additionally, we observed high exudation of pyruvate coming from the pentose phosphate and Entner-Doudoroff pathways. Both are alternative routes coming out of carbon fixation (Waldbauer et al., 2012; Chen et al., 2016). Together with a low biomass value, this phenotype might indicate a scenario of exudation due to overflow metabolism.

**Figure 4:**
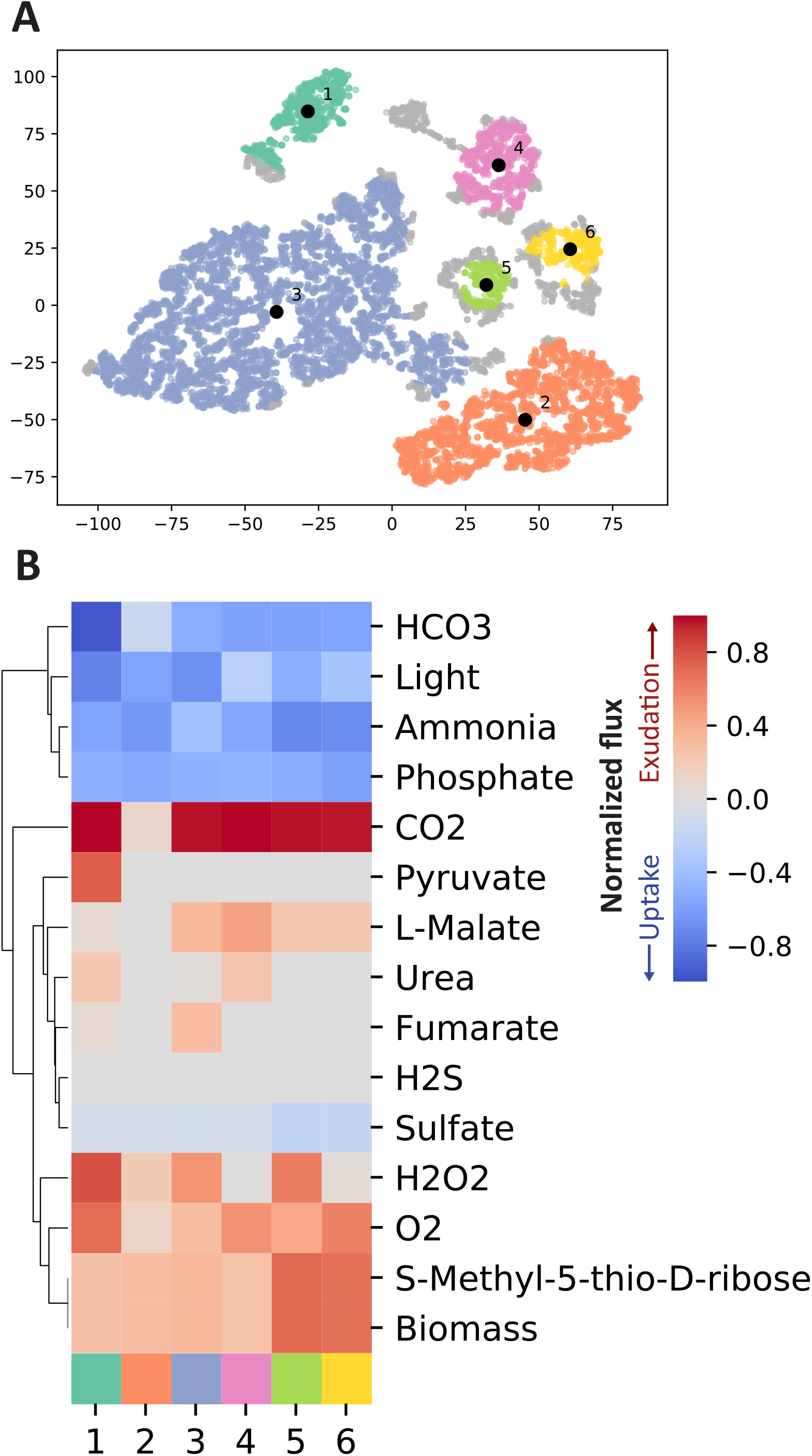
T-SNE clustering identifies typical phenotypes from the pFBA results from the random samples. **A)** The random samples are reduced into two dimensions with t-SNE. We have subsequently used HDBSCAN to identify 6 disjoint clusters which represents different phenotypes. **B)** For each of the 6 clusters the mean uptake or exudation across all samples within the respective cluster is shown. Only exchange reactions with an absolute flux above 1e-3 mmol gDW^−1^ h^−1^ in any of the random samples are included.

The two largest clusters (number 2 and 3), tie together high and low light, carbon and nitrogen uptake rates, and different exudation patterns. Interestingly, phenotype 3 (high light) showed exudation of fumarate and malate while phenotype 2 (low light) did not. Recent work suggested that, in high light conditions, fumarate is generated through oxaloacetate and malate creating a broken acyclic form of the TCA cycle, while in the dark, fluxes are diverted into forming the cyclic form of it. This low light form of the TCA cycle is then active and works towards energy generation (Xiong et al., 2017). Similarly, we observed two forms of the TCA cycle in the high and low light phenotypes (2 and 3 respectively) with a difference in the direction of one reaction (KEGG R00342, Figure S3). Phenotype 2, describing low-light conditions, showed the L-Malate/oxaloacetate balance to shift in favor of oxaloacetate, completing the route towards 2-Oxoglutarate, a key metabolite known to act as a starvation signal and modulator of the C/N balance in cyanobacteria (Domínguez-Martín et al., 2018; Zhang et al., 2018), and subsequently into energy generation. On the other hand, Phenotype 3, describing high light conditions, showed the L-Malate/oxaloacetate balance to shift in favor of L-Malate and away from the formation of 2-oxoglutarate. In both phenotypes fumarate is converted to L-Malate. While in Phenotype 2 it is fed into a semi-cyclic form of the TCA cycle, fumarate is partly exuded and partly converted to L-malate in phenotype 3, in agreement with overflow metabolism.

We observed a similar TCA cycle flux distribution in phenotype 4 leading to high exudation of L-Malate. Interestingly, Phenotype 1 and 4 are comparable in all key nutrients except light (High in phenotype 1 and low in phenotype 4). As a result of an in-depth flux distribution analysis, we observed a reaction direction change in UDP-glucose:NAD+ 6-oxidoreductase [R00286, EC 1.1.1.22, PMM1261] between the two phenotypes. In phenotype 4 this reaction shifted towards the creation of UDP-glucose, a precursor for the production of glycogen (Due to the high stringency of this analysis we did not observe the direct formation of glycogen). In phenotype 1, this reaction favored the formation of UDP-glucuronate which in turn was diverted into the formation of amino sugars. These phenotypes may correlate to the 12:00 (phenotype 1) and 16:00 (phenotype 4) scenarios described in (Szul et al., 2019). Finally, Phenotypes 5 and 6 may represent a high-light nutrient-rich environment resulting in a high biomass value.

### Nutrient uptake rates modulate exudation of organic compounds

The use of genome-scale metabolic models captures a comprehensive picture of the metabolic processes taking place in the cell, including those that lead to metabolite exudation. From the random sampling of environmental conditions, we identified conditions in which organic acids must be exuded. This was noticeable by a non-zero lower bound of the FVA region (Figure 3). Interestingly, organic acids were more likely to be exuded when the growth became limited by phosphate or nitrogen. Since *Prochlorococcus* is known to thrive in oligotrophic ocean gyres where nitrogen or phosphate is limited (Partensky et al., 1999; Flombaum et al., 2013b), this represents a likely natural phenotype, and as such, supports previous findings (Szul et al., 2019). Costly metabolites, essential for cell survival and growth, such as amino acids, nucleobases and nucleotides, tend to be exuded in nitrogen and carbon rich conditions and might be a result of overflow metabolism (Cano et al., 2018; Pacheco et al., 2018). To explore this phenomenon in further detail, we looked into exudation patterns of specific metabolites as a function of key nutrient limitations (Figure 5). Of the environmental factors, the uptake of nitrogen (ammonium) is a decisive factor differentiating between exudation of organic acids or amino acids. While it is positively correlated with the exudation of nitrogen-rich compounds such as amino acids, it is negatively correlated with exudation of organic acids and glycogen. Additionally, glycogen formation is positively correlated with the exudation of malate, citrate, fumarate and succinate, which are most of the TCA cycle constituents. This is in line with previous findings suggesting the re-direction of carbon metabolism towards the formation of macromolecules (including glycogen) in nitrogen limiting conditions (Forchhammer and Selim, 2019; Szul et al., 2019). Thus, our reconstruction captured known possible aspects of the carbon/nitrogen balance in *Prochlorococcus*.

**Figure 5:**
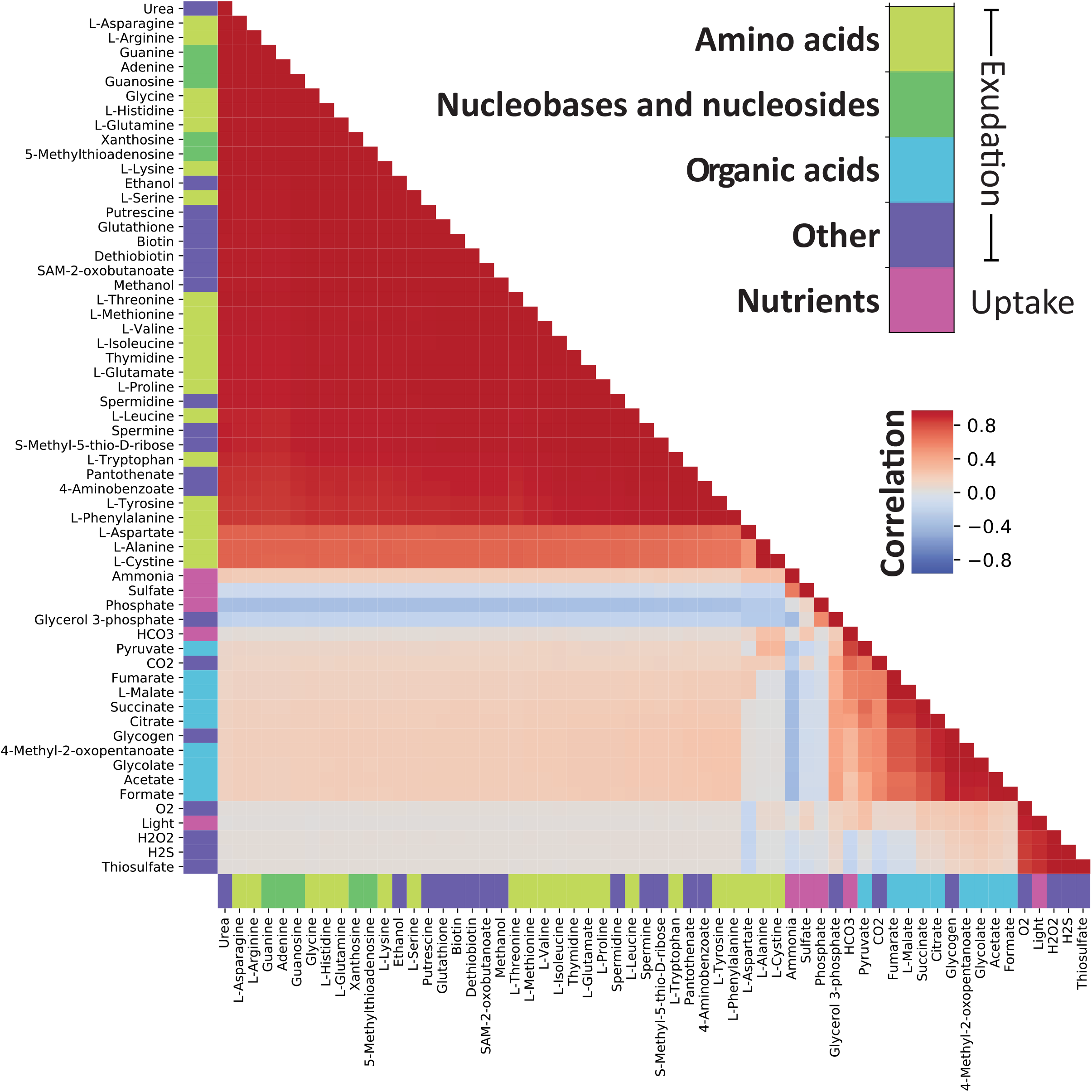
**Correlation between maximal FVA values** demonstrate which compounds that can be secreted in similar environmental conditions. There are two clearly correlated groups of compounds: The first group comprises nucleobases, nucleotides and amino acids while the second contains organic acids, glycogen and glycerol 3-phosphate. From the correlation of each of these groups with the uptake bounds on ammonium and phosphate we observe that these factors determine which of the groups that can be secreted. Note that the environmental constraints have been converted to positive values prior to calculating the correlation.

Finally, we observed a general pattern of strong positive correlations between amino acids, nucleobases, nucleosides, as well as a range of other compounds. In an interesting deviation from this general pattern, L-Aspartate showed a decreased correlation with other exudates. L-Aspartate, together with its role in protein nucleic acid biosynthesis, can serve as a precursor for nitrogen storage metabolites such as polyamines (Szul et al., 2019). Indeed, we observed a slightly stronger correlation between L-Aspartate and the uptake of nitrogen compared to other amino acids. Finally, In contrast to other amino acids, L-Aspartate is negatively correlated with light uptake and hydrogen peroxide exudation. Hydrogen peroxide is produced from L-Aspartate and oxygen by L-aspartate oxidase [R00481, EC 1.4.3.16, PMM0100]. L-amino acid oxidases have been previously described in cyanobacteria and have been related to the use of amino acids as carbon sources (Campillo-Brocal et al., 2015). The production of hydrogen peroxide is also strongly correlated with light, a result consistent with the expectation that reactive oxygen species are created during photosynthesis.

### Dynamic allocation of carbon storage

Nutrient and light limitations are well-known modulators of carbon storage in *Prochlorococcus* (Zinser et al., 2009; Szul et al., 2019). Recent work has suggested the storage of carbon to be one of the major metabolic tasks during the day-night cycle (Cano et al., 2018; Szul et al., 2019; Shinde et al., 2020). Sampling 10,000 environmental conditions showed that glycogen production decreased at lower light intensities and was less saturated at higher rates (Figure 3). To explore other time-modulated trade-offs and trends related to carbon storage, we performed *in silico* dynamic FBA diel-cycle simulations using the Computational of Microbial Ecosystems in Time and Space (COMETS) platform (Mahadevan et al. 2002; Harcombe et al. 2014). COMETS relies on kinetic information such as K_m_ and V_max_ to simulate the spatial growth and exudation patterns of microbes in a simulated discretized time course. To improve the accuracy and biological relevance of our simulations we used kinetic constants either obtained from experimental measurements reported in the literature (Krumhardt et al., 2013; Hopkinson et al., 2014) (Table S3) or from fitting model simulations to measured growth and depletion of ammonium rates (Grossowicz et al., 2017). We found K_m_ and V_max_ values of 0.39 mM and 0.9 mmol gDW^−1^ h^−1^ for the uptake of ammonium to best fit the experimental data (Grossowicz et al., 2017) (Figure S1). Surprisingly, the estimated K_m_ value is 3 orders of magnitude larger than previous estimates (Grossowicz et al., 2017), indicating some uncertainty in this estimate that may be affected by the choice of method for calculating this value.

Since the tight coupling between carbon and nitrogen metabolism in cyanobacteria is known to influence carbon allocation and storage (Zhang et al., 2018; Szul et al., 2019), it was chosen as a case study. As such, we focused in more detail on the dynamic changes in metabolism in nitrogen-abundant and nitrogen-poor media, as previously defined (Grossowicz et al., 2017). Specifically, we set out to explore glycogen production and consumption with COMETS in these conditions (Figure 6). We did not observe glycogen storage in nitrogen-abundant simulations. One explanation for this may arise from the limitations of the platform. The modelling framework is based on linear programming to optimize an ordered set of assumed cellular objectives, glycogen is only stored when there is excess energy and carbon available, which is the case when the growth is nitrogen limited. Another limitation that might affect glycogen storage in these simulations might involve other types regulatory mechanisms not usually accounted for in dynamic FBA. The addition of regulatory layers, such as proteomics data (Reimers et al., 2017), or more specifically tailored objective functions could lead to smaller but nonzero generation of glycogen also during nitrogen-rich conditions.

**Figure 6:**
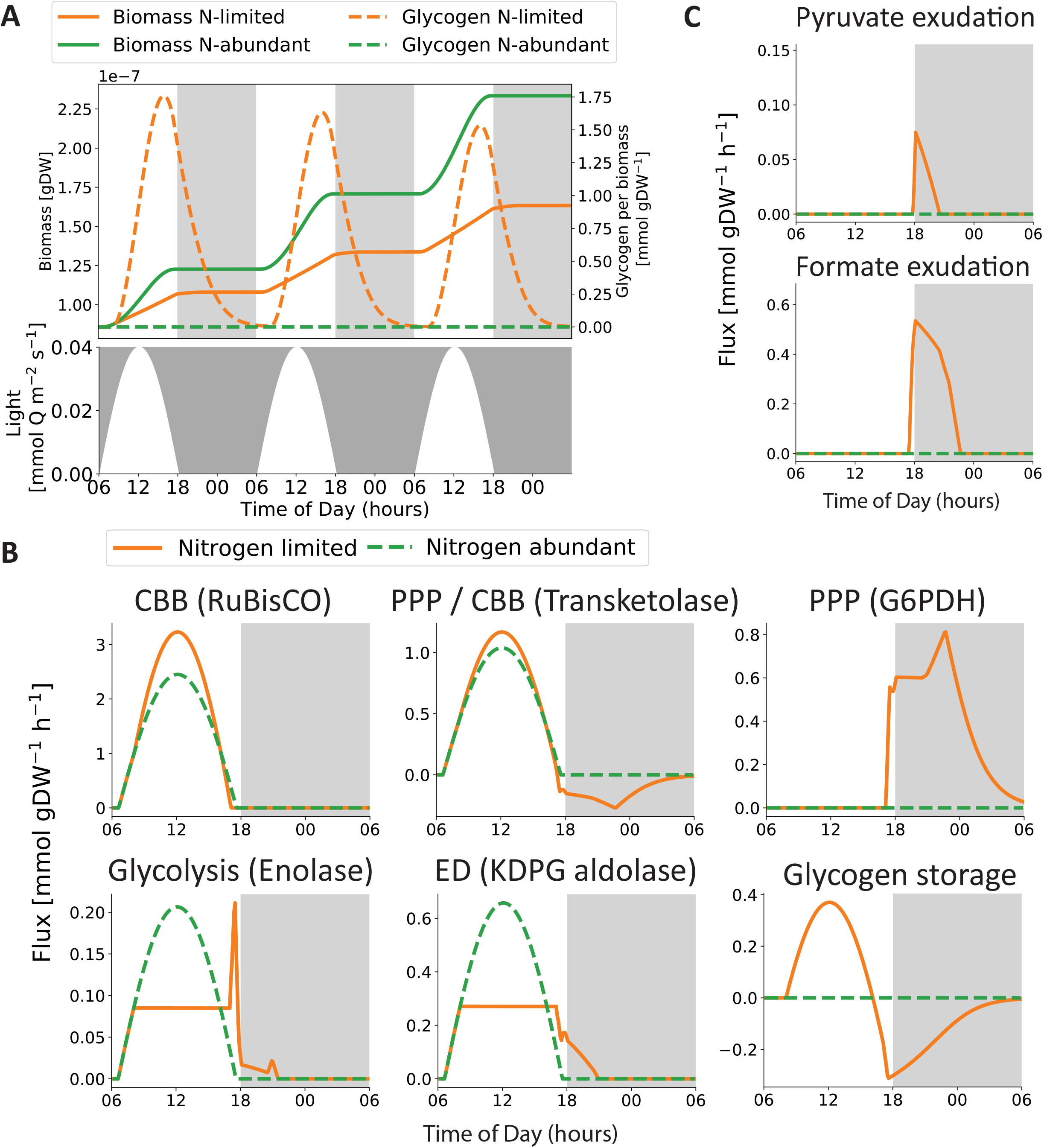
Insight into metabolic rearrangements during the diel cycle. **A)** Light irradiance, biomass and glycogen storage throughout the diel cycle. We observe that the largest accumulated growth is found in the nitrogen-abundant condition (green), but glycogen is only predicted to be stored in the nitrogen-poor condition (orange). **B)** The flux distributions shifts when the metabolism switches from photosynthesis to glycogen catabolism, displayed by five reactions representing the Calvin Cycle (CBB), Glycogen metabolism, lower part of glycolysis, The Entner-Doudoroff (ED) and the Penthose Phosphate Pathway (PPP). **C)** iSO595 predicts that the depletion of glycogen is accompanied by exudation of fumarate and pyruvate.

In agreement with previous work (Szul et al., 2019), under nitrogen-limiting conditions, glycogen accumulates throughout the day and is subsequently used to support respiration and growth during the night (Figure 6A). However, the predicted glycogen storage is not sufficient to support neither growth nor cellular maintenance during the night and may explain the reported increased death rate during night time (Zinser et al., 2009; Ribalet et al., 2015). Interestingly, the model predicts consumption of glycogen during dusk to increase growth when photosynthesis is declining (Figure 6), closely resembling observations in *Synechococcus*, in particular for the Δ*kaiC* mutant with a dysfunctional circadian clock (Diamond et al., 2015).

The switch from photosynthesis at daytime to glycogen consumption at nighttime is reflected in the metabolic shifts observed in key pathways (Figure 6B). Interestingly, we observed higher fluxes through the Calvin cycle in nitrogen-poor conditions. This difference may be caused by the increased ATP demand necessary to support higher growth rates in nitrogen-abundant conditions. Additionally, our simulations predicted that the use of the Entner-Doudoroff pathway during photosynthesis creates precursor metabolites for growth during light hours, and a shift to the Pentose Phosphate Pathway (PPP) during nighttime. This trend might occur as an alternative for generating NADPH (Figure S4). Upregulation of the PPP enzymes during dusk and the first half of the night time was also observed in the proteome of *Prochlorococcus* (Waldbauer et al., 2012). Several enzymatic transformations participate in both the Calvin cycle and the PPP, although in opposite directions (Waldbauer et al., 2012). These transformations were captured in our simulations, specifically as demonstrated by transketolase (Figure 6B). Additionally, the consumption of glycogen during nighttime leads to exudation of pyruvate and formate (Figure 6C). This prediction is supported by previous observations; formate is exuded when phosphonates are metabolized in the closely related strain *Prochlorococcus marinus* MIT9301 (Sosa et al., 2019) and pyruvate is exuded when fixed carbon is consumed in *Synechococcus* elongatus PCC 7942 and *Synechocystis* sp. PCC 6803 (Carrieri et al., 2012; Benson et al., 2016).

The shift from photosynthesis and carbon fixation to glycogen catabolism is also associated with a switch in production and consumption of energetic cofactors (Figure S3). Generation of ATP is performed concomitantly by ATP synthase in both the thylakoid membrane and the periplasmic membrane during photosynthesis. The periplasmic ATP synthase is first driven by reduced cofactors (NADPH) generated by the electron transport chain in the light-dependent part of photosynthesis (Figure S3). ATP is consumed by two separate processes: growth- and maintenance-associated reactions reach a threshold once growth is limited by the nitrogen abundance, while the recycling of precursors for the Calvin cycle follows the shape of light absorption throughout the day. In agreement with previous work (Park and Choi, 2017), our model predicted higher rates of NADPH production than NADH.

Next, we explored the ability of our model to dynamically capture biologically relevant phenotypes by performing dynamic FBA simulations of knock-out mutants in *Prochlorococcus*, focusing on two gene deletions disrupting different parts of glycogen metabolism. Δ*glgC* breaks synthesis of ADP-glucose and thus the storage of glycogen and Δ*gnd*, knocking out 6-phosphogluconate dehydrogenase, a key reaction in the Pentose Phosphate pathway found to fuel the Calvin cycle with precursor metabolites during the onset of photosynthesis (Shinde et al., 2020). Our dynamic FBA simulations in COMETS (Figure S5) showed similar growth between Δ*gnd* and the wild type and slightly lower growth for Δ*glgC*. We set out to compare these observations with available experimental data. Since genetic tools for the modification of *Prochlorococcus* are still lacking (Laurenceau et al., 2020), we chose data from the closely related cyanobacteria *Synechococcus* as recent work described the impact of Δ*glgC* and Δ*gnd* on its growth during diel cycles (Shinde et al., 2020). Indeed, we found very good agreement between measured and predicted growth for both the wild-type and Δ*glgC* mutant where glycogen storage is disrupted (Shinde et al., 2020) (Figure S5). One of the notable limitations of dynamic FBA is the ability to quantify intermediates and precursor pools that might drive the initiation of a pathway. This comes mainly from the assumption of a quasi steady-state of intracellular metabolite pools at each time point. Although the comparison is strictly qualitative and concerns strains with known differences (Mary et al., 2004), these findings demonstrated the ability of our reconstruction to capture metabolic trends in response to genetic perturbations, indicating that *i*SO595 will be a valuable tool in future research of *Prochloroccus*. Overall, our dynamic simulations display biological and physiological behaviors that are consistent with expectations, and at the same time provide valuable insight into the putative internal metabolic processes that might modulate the *Prochlorococcus* growth under environmental and genome-induced constraints.

## Conclusions

Our study provides a detailed systematic view of the underlying metabolic trends modulating carbon storage and exudation in *Prochlorococcus. Prochlorococcus* is known to interact with other bacteria in its surroundings (Sher et al., 2011; Aharonovich and Sher, 2016; Biller et al., 2018; Hennon et al., 2018). It is currently impossible to predict the fluxes of organic matter (or of the myriad metabolites comprising it, such as amino acids, sugars and organic acids) between phytoplankton and bacteria. Yet, quantifying such fluxes and predicting them from genomic surveys, as shown here, serves a number of roles: (1) It can provide experimentally testable and mechanistic hypotheses on inter-microbial exchanges and competition, (2) It has the potential to increase knowledge about the specific metabolites that may mediate these interactions; and (3) It would enable the construction of improved models of biogeochemical cycles which consider the diverse and powerful metabolic capabilities of the ocean microbiome.

Genome-scale metabolic-network reconstructions are a powerful tool, but not without limitations. With the development of an innovative semi-automated approach (the ReFill algorithm), we were able to increase existing knowledge in the reconstruction by up to 25%. In contrast to other standard gap filling approaches (Henry et al., 2010; Thiele et al., 2014), or plain addition of newly found reactions, ReFill has the specific capability to add individual reactions through a recursive algorithm that guarantees complete connectivity to the existing network. However, this approach employs high stringency and thus adds limited amounts of knowledge. Considering that *Prochlorococcus* strains have some of the smallest known genomes among free-living organisms, a 25% increase in knowledge serves as a significant improvement in the predictive capacity of the model. However, reconstruction of high-quality genome-scale metabolic models is an iterative process, where new data, knowledge, and scope creates a demand and possibility for further model improvement. One example of this possibility is the CO_2_ concentrating mechanisms in *Prochlorococcus*. This mechanism is known to be sustained by proton and ion gradients across the cell membrane at an energetic cost (Hopkinson et al., 2014; Burnap et al., 2015). However, the comprehensive knowledge and annotation of ion transporters necessary to model this mechanism are lacking, and therefore, not included in iSO595. With the advancement of data collection and annotation tools, together with using ReFill or similar algorithms, metabolic knowledge can be easily added to such reconstructions and improve their predictive abilities and mimicry of biological and physiological processes. Our findings contribute to a growing body of work on the underlying metabolic mechanisms modulating the metabolic success of *Prochlorococcus*. The approaches shown here provide systematic insights corroborated in recent and well-known works and provide strong foundations for future studies of *Prochlorococcus* metabolism with particular interest in its interaction with other microorganisms and the effects of these on community composition and larger biogeochemical cycles.

## Acknowledgments

This work is available as preprint on bioRxiv in https://doi.org/10.1101/2020.07.20.211680 (Ofaim et al., 2020)

## Author contributions

DSh and DSe designed the study. SO, SS and DSe developed the computational models and performed the computational analyses. SO and SS wrote a first version of the manuscript. DSe, DSh and EA, oversaw the project and contributed to the final version of this manuscript. All authors have read and approved the final version of the manuscript.

## Supplemental material

**Figure S1: Estimation of kinetic parameters for the uptake of ammonium in *Prochlorococcus*. A)** All combinations of Km and Vmax along the red trajectory matches the observed gross growth rate of 0.5 d^−1^ (Grossowicz et al., 2017). However, when we compare the dynamics of cell density (B) and ammonium concentration (C) we find that the best overall prediction is achieved using Km = 0.39 mM and Vmax = 0.9 mmol gDW^−1^ h^−1^ (marked by an orange dot in **A).**

**Figure S2: ATP availability influence modulates the trade-off between glycogen storage and growth.** Phenotypic phase planes (Edwards et al., 2002) illustrate the combined effect of glycogen storage and bicarbonate uptake on the maximal growth rate. Compared to the base model (A), we observe how that the trade-off is strongly affected by modulated ATP availability, either from an artificial reaction providing extra ATP **(B)** or by increasing **(C)** or decreasing **(D)** the amount of available light. Increasing ATP allows more glycogen storage without reducing the growth rate.

**Figure S3: TCA cycle flux diagram differences between the most common phenotypes.** Flux diagrams of the TCA cycle in the most common phenotypes 2 (colored orange) and 3 (colored blue). Reactions are denoted by KEGG reaction ids. Reaction colors correspond to cluster colors presented in figure 4.

**Figure S4: The transition from daytime to nighttime is associated with a drastic change in the production and consumption of the energy-carrying cofactors. The figure panels show the major sources (left) and drains (right) of the cofactors ATP, NAPDH and NADH.** The legend shows the reaction IDs used in iSO595. **A)** ATP is produced by both the thylakoid (R00086th) and the periplasmic (R00086p) ATP synthase during daytime, but mostly by the periplasmic ATP synthase (respiration) during nighttime. ATP is consumed by reactions associated with growth (BIOMASS and BProtein), cellular maintenance (Maintenance) and storage of glycogen (R00948) in addition to reactions recycling precursors for the Calvin cycle (R01512 and R01523) and acetyl-CoA carboxylase (R00742). **B)** NADPH is produced by ferredoxin reductase (fdr) during daytime and by the pentose phosphate pathway (R02736 and R01528) during nighttime. The NADPH is either used to drive the proton gradient across the periplasmic membrane (NADPHDHp) or in Gluconeogenesis (R01063) to refuel the Calvin cycle during photosynthesis. **C)** NADH production is correlated with the growth rate and dominated by pyruvate dehydrogenase (R00209) during daytime and the glycine cleavage system (R01221) during nighttime. NADH is consumed by 6-phosphogluconate dehydrogenase (reverse, R10221) during daytime, NADH transhydrogenase (R00112) solely during dusk and concomitant with methylenetetrahydrofolate reductase (R07168) during nighttime.

**Figure S5: Predicted growth curves show good agreement in a qualitative comparison with experimental growth of *Synechococcus*.** To compare growth data we have overlaid growth curves predicted for the wild-type, the Δ*glgC*-mutant and the Δ*gnd*-mutant of *Prochlorococcus* with experimental OD measurements of *Synechococcus elongatus PCC 794* (Shinde et al., 2020). We find a very good agreement for the wild-type and Δ*glgC*-mutant, but not for the Δ*gnd*-mutant. The lower panel shows how the model predicts the allocation and consumption of each of the three strains.

**Table S1:** List of reactions added to iJC568 to form iSO595.

**Table S2:** List of reactions from iJC568 that are modified in iSO595.

**Table S3:** Parameter ranges used in the sampling of nutrient environments.

**Table S4:** List of blocked exchange reactions prior to sampling of nutrient environments.

**Table S5:** Parameter values used to run dynamic FBA in COMETS.

**Supplemental material 1:** Results from the BLAST-search used to identify 6PG-dehydratase (EC: 4.2.1.12) encoded by PMM0774 in *P. marinus* MED4.

**Supplemental material 2:** memote snapshot report of *i*SO595.

