## Supplementary figures and images for "Dynamic allocation of carbon storage and nutrient-dependent exudation in a revised genome-scale model of *Prochlorococcus*"

### Supplemental figures

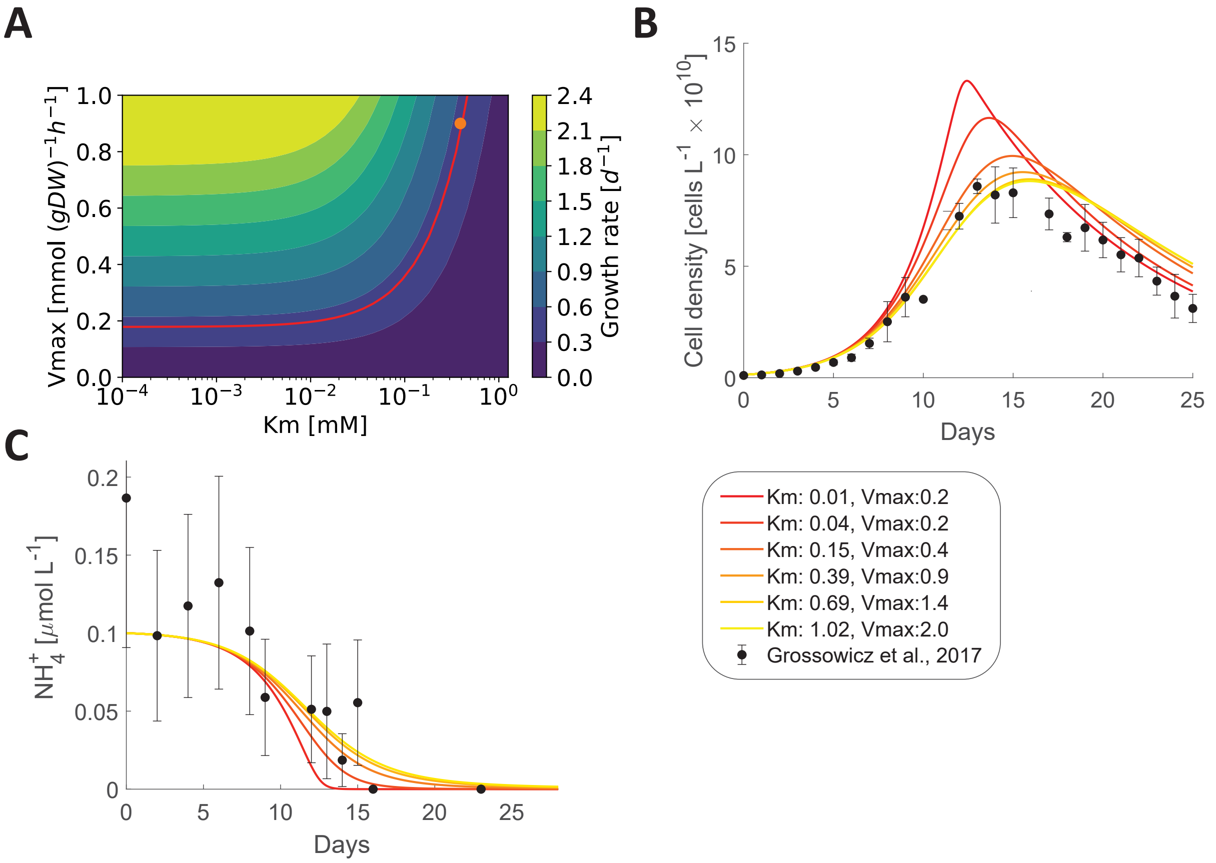

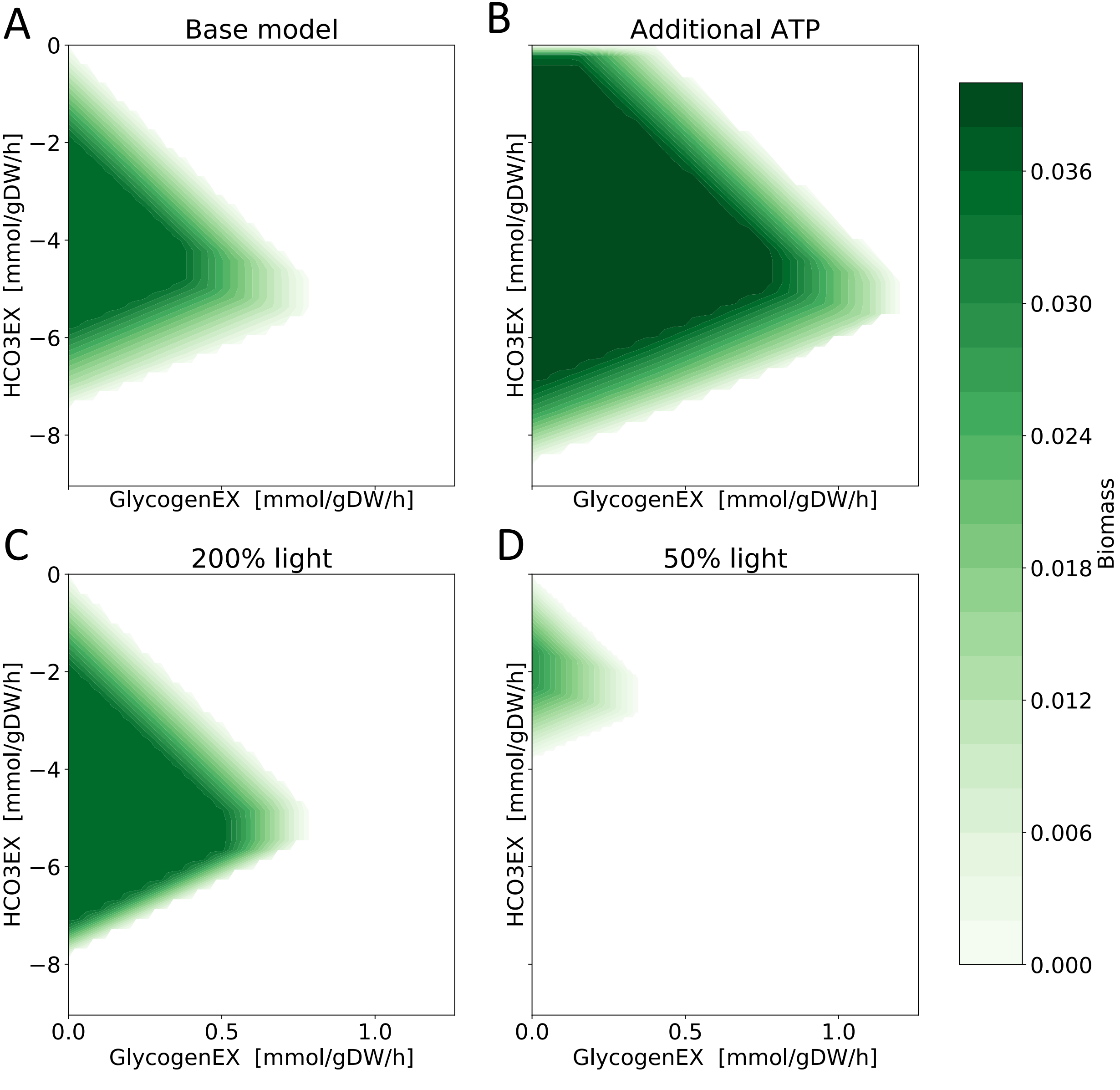

#### Phenotype 2

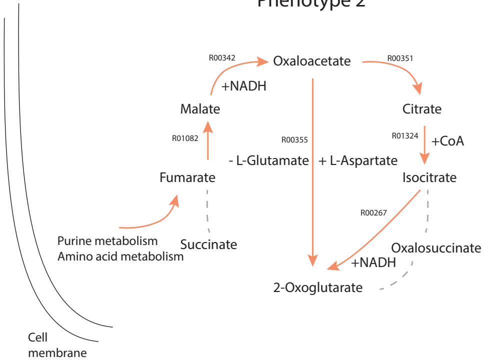


#### Phenotype 3

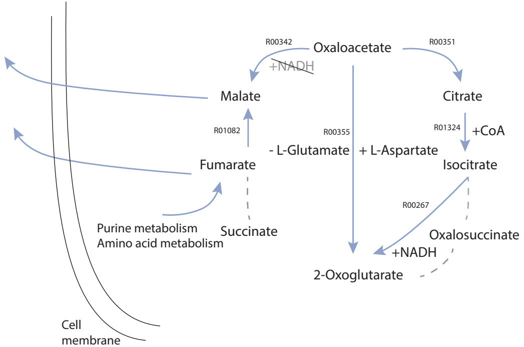

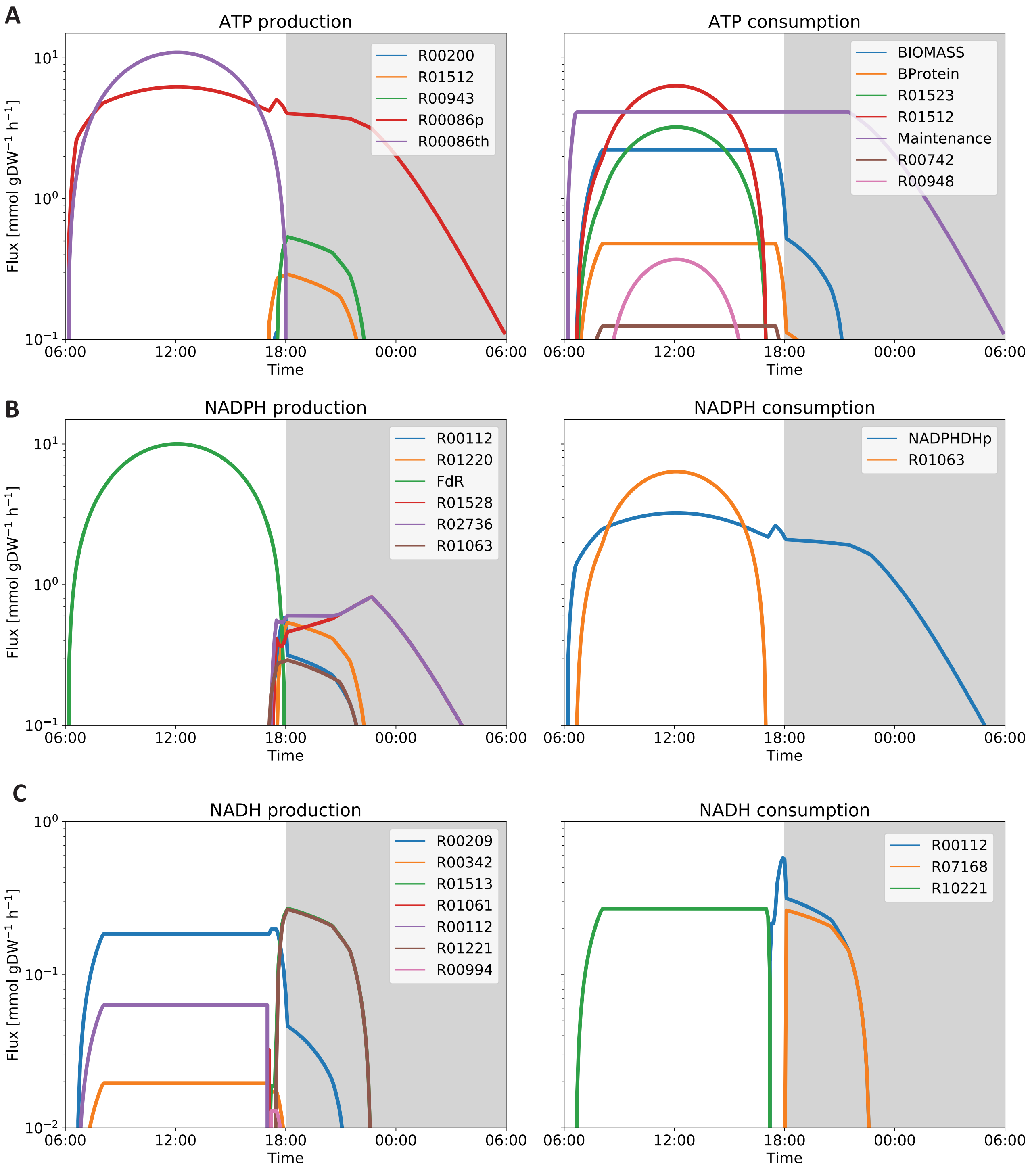

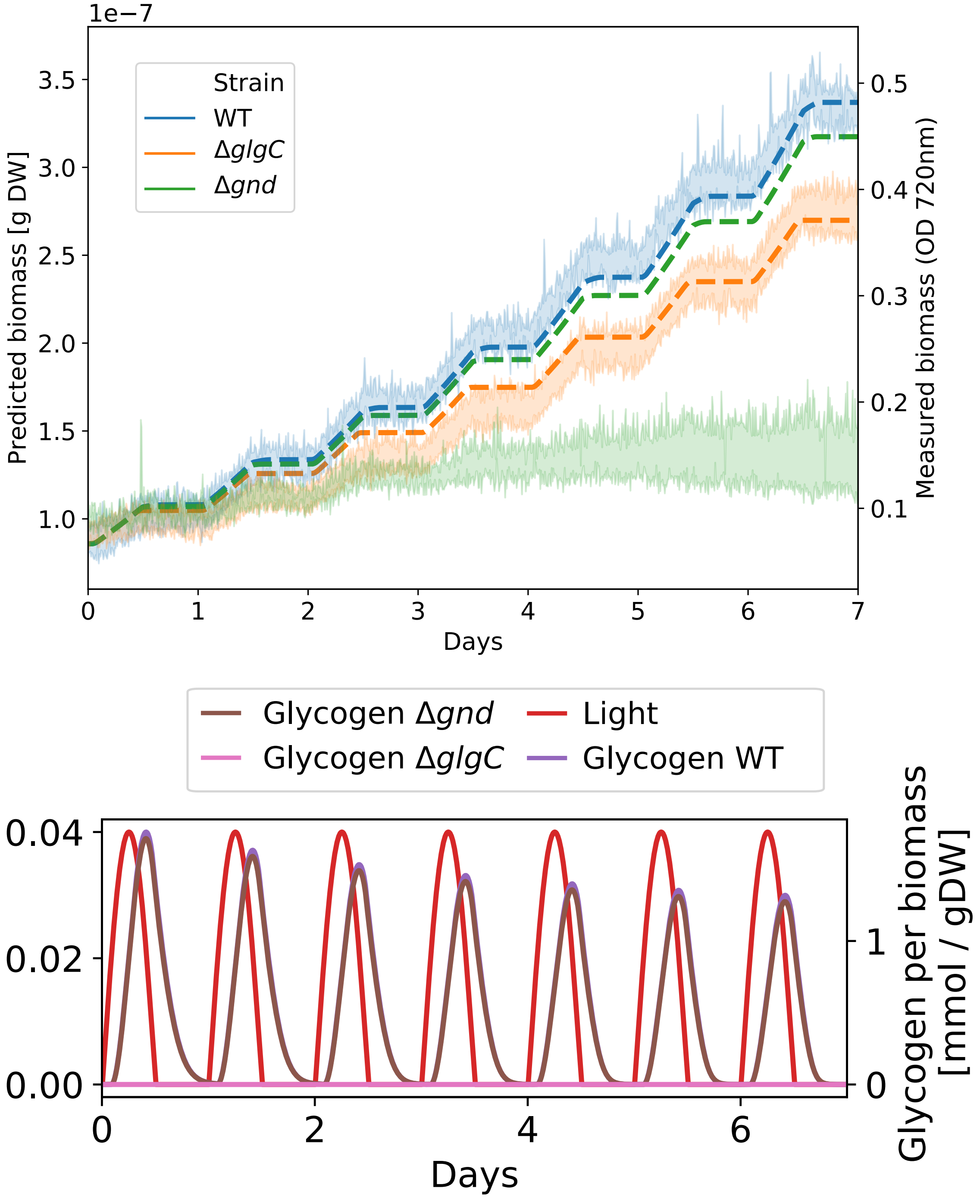
