## Supplemental material 1 for "Dynamic allocation of carbon storage and nutrient-dependent exudation in a revised genome-scale model of *Prochlorococcus*"

BLASTP 2.2.29+

Reference: Stephen F. Altschul, Thomas L. Madden, Alejandro A. Schaffer, Jinghui Zhang, Zheng Zhang, Webb Miller, and David J. Lipman (1997), "Gapped BLAST and PSI-BLAST: a new generation of protein database search programs", Nucleic Acids Res. 25:3389-3402.

Reference for composition-based statistics: Alejandro A. Schaffer, L. Aravind, Thomas L. Madden, Sergei Shavirin, John L. Spouge, Yuri I. Wolf, Eugene V. Koonin, and Stephen F. Altschul (2001), "Improving the accuracy of PSI-BLAST protein database searches with composition-based statistics and other refinements", Nucleic Acids Res. 29:2994-3005.

Database: /bio/db/fasta/genes/T00004.pep
3,564 sequences; 1,136,958 total letters

Query= pmm:PMM0774 K01687 dihydroxy-acid dehydratase [EC:4.2.1.9] |
(GenBank) ilvD; Dihydroxy-acid dehydratase (A)

Length=559

Sequences producing significant alignments:

Top 5 | Clear | Select operation | Exec

| | Sequence | Description | K number | Score (Bits) | E Value |
|---|---|---|---|---|---|
| ☑ | syn:slr0452 | ilvD; dihydroxyacid dehydratase | K01687 | 838 | 0.0 |
| ☑ | syn:sll0154 | hypothetical 35.6 kD protein | | 29.6 | 0.47 |
| ☐ | syn:slr1219 | ureE; urease accessory protein E | K03187 | 26.6 | 2.5 |
| ☐ | syn:sll0180 | unknown protein | | 26.9 | 3.2 |
| ☐ | syn:slr0326 | unknown protein | | 26.2 | 3.6 |
| ☐ | syn:sll1866 | hypothetical protein | K07566 | 25.4 | 7.2 |
| ☐ | syn:sll1513 | ccsA; c-type cytochrome synthesis protein | | 25.8 | 7.8 |

>syn:slr0452 ilvD; dihydroxyacid dehydratase ↑ Top
Length=561

```
 Score =   838 bits (2166),  Expect = 0.0, Method: Compositional matrix adjust.
 Identities = 391/557 (70%), Positives = 474/557 (85%), Gaps = 0/557 (0%)

Query  2    NKLRSSAITQGVQRSPNRSMLRAVGFSDEDFTKPIIGVANGFSTITPCNMGLNKLALKAE  61
            N  RS  ITQG QRSPNR+MLRAVGF D+DFTKPI+G+ANG+STITPCNMG+N LAL+AE
Sbjct  3    NNPRSQVITQGTQRSPNRAMLRAVGFGDDDFTKPIVGIANGYSTITPCNMGINDLALRAE  62

Query  62   ESIREAGGMPQMFGTITVSDGISMGTEGMKYSLVSREVIADSIETACNAQSMDGVLAIGG  121
              +R AG MPQ+FGTIT+SDGISMGTEGMKYSLVSREVIADSIET CN Q MDGVLAIGG
Sbjct  63   AGLRTAGAMPQLFGTITISDGISMGTEGMKYSLVSREVIADSIETVCNGQRMDGVLAIGG  122

Query  122  CDKNMPGAMIAIARMNIPSIFIYGGTIKPGKLNGEDLTVVSAFEAVGQLTSGKINEKRLI  181
            CDKNMPGAMIA+AR+NIPSIF+YGGTIKPG   GEDLTVVSAFEAVGQ ++GKI+E+ L
Sbjct  123  CDKNMPGAMIAMARLNIPSIFVYGGTIKPGHYAGEDLTVVSAFEAVGQYSAGKIDEETLY  182

Query  182  EVEKNCIPGAGSCGGMFTANTMSAVIEVLGLSLPYSSTMAAEDYEKEVSAEKSAEILVDA  241
             +E+N  PGAGSCGGMFTANTMS+  E +G+SLPYSSTMAA D EK  S E+SA++LV+A
Sbjct  183  GIERNACPGAGSCGGMFTANTMSSAFEAMGMSLPYSSTMAAVDGEKADSTEESAKVLVEA  242

Query  242  IRKDIRPLTLMTKESFENAITVIMAIGGSTNAVLHILAIANTAGIDINIDDFERIRQKVP  301
            I+K I P  ++T+++FENAI VIMA+GGSTNAVLH+LAIANT G+ +++DDFE IR KVP
Sbjct  243  IKKQILPSQILTRKAFENAIAVIMAVGGSTNAVLHLLAIANTIGVPLSLDDFETIRHKVP  302

Query  302  VICDLKPSGKYVTVDLHKAGGIPQVMKILLNTGLIHGNCRNIEGKTVVESLKDIPVKPPE  361
            V+CDLKPSGKYVT +LH AGGIPQVMKILL  G++HG+   I G+T+ E L DIP +PP
Sbjct  303  VLCDLKPSGKYVTTNLHAAGGIPQVMKILLVNGILHGDALTITGQTIAEVLADIPDQPPA  362

Query  362  NQDVIRDIDNPLYKKGHLAILKGNLASEGCVAKISGIKNPVLKGPARIFESEEDCLKSIL  421
             QDVI   D+P+Y++GHLA+LKGNLA+EG VAKISG+K PV+ GPA++FESEEDCL++IL
Sbjct  363  GQDVIHSWDDPVYQEGHLAVLKGNLATEGSVAKISGVKKPVITGPAKVFESEEDCLEAIL  422

Query  422  NNDIKAGNVVVIRNEGPVGGPGMREMLAPTSAIVGQGLGEKVALITDGRFSGGTYGLVVG  481
               I+AG+VVV+R EGP GGPGMREMLAPTSAI+G GLG+ V LITDGRFSGGTYGLVVG
Sbjct  423  AGKIQAGDVVVVRYEGPKGGPGMREMLAPTSAIIGAGLGDSVGLITDGRFSGGTYGLVVG  482

Query  482  HIAPEAAVGGNIALIKEGDLITVDATNQLIEVELSDEELEMRRINWEKPSKKYKKGVLSK  541
            H+APEA VGG IAL++EGD IT+DA  +L+++ +S+EEL  RR  W  P  +Y +G+L+K
Sbjct  483  HVAPEAYVGGAIALVQEGDQITIDAGKRLLQLNISEEELAQRRAQWTPPQPRYPRGILAK  542

Query  542  YSRIVSTSSLGAVTDLE  558
```

```
            Y+++VS+SSLGAVTD++
Sbjct  543  YAKLVSSSSLGAVTDID  559


>syn:sll0154 hypothetical 35.6 kD protein                                    ↑ Top
Length=462

 Score = 29.6 bits (65),  Expect = 0.47, Method: Compositional matrix adjust.
 Identities = 15/57 (26%), Positives = 31/57 (54%), Gaps = 2/57 (4%)

Query  126  MPGAMIAIARMNIPSIFIYGGTIKPGKLNGEDLTVVSAFEAVGQLTSGKINEKRLIE  182
            +P    AIA+  +P I+I     +PG+   ++ TV     A+ Q++S ++ +  L++
Sbjct  341  IPEIQTAIAKAKVPRIYICNVMTQPGE--TDNYTVSDHLTAIDQVSSARLYDAVLVQ  395


>syn:slr1219 ureE; urease accessory protein E                                ↑ Top
Length=142

 Score = 26.6 bits (57),  Expect = 2.5, Method: Composition-based stats.
 Identities = 16/42 (38%), Positives = 24/42 (57%), Gaps = 7/42 (17%)

Query  139  PSIFIY---GGTIKPGKL----NGEDLTVVSAFEAVGQLTSG  173
            PS+FI    G  ++PG       GE +T+++A E +  LTSG
Sbjct  39   PSLFIQLPRGSFLRPGDCLGSPTGETITILAADEPLLHLTSG  80


>syn:sll0180 unknown protein                                                 ↑ Top
Length=501

 Score = 26.9 bits (58),  Expect = 3.2, Method: Compositional matrix adjust.
 Identities = 17/55 (31%), Positives = 26/55 (47%), Gaps = 1/55 (2%)

Query  423  NDIKAG-NVVVIRNEGPVGGPGMREMLAPTSAIVGQGLGEKVALITDGRFSGGTY  476
            ND++ G  V V+R++G  G  G    ++PT+    Q +  KV    DG      Y
Sbjct  333  NDLRLGLPVEVVRSDGQAGEVGRISFISPTANRNDQSILAKVIFKNDGSLRNNQY  387


>syn:slr0326 unknown protein                                                 ↑ Top
Length=147

 Score = 26.2 bits (56),  Expect = 3.6, Method: Composition-based stats.
 Identities = 14/39 (36%), Positives = 24/39 (62%), Gaps = 2/39 (5%)

Query  292  DFERIRQKVPVICDLKPSGKYVTVDLHKAGGIPQVMKIL  330
            + +   QKV VIC+L+ +GK  T++ ++   I Q+ K L
Sbjct  89   NLQEFAQKVSVICNLETAGKIETMEAYER--IKQLWKSL  125


>syn:sll1866 hypothetical protein                                            ↑ Top
Length=199

 Score = 25.4 bits (54),  Expect = 7.2, Method: Compositional matrix adjust.
 Identities = 11/29 (38%), Positives = 17/29 (59%), Gaps = 0/29 (0%)

Query  327  MKILLNTGLIHGNCRNIEGKTVVESLKDI  355
            ++ILL TG +     N+ G+  +E L DI
Sbjct  120  LEILLQTGPLATTSANLSGQPPLEKLADI  148


>syn:sll1513 ccsA; c-type cytochrome synthesis protein                       ↑ Top
Length=334

 Score = 25.8 bits (55),  Expect = 7.8, Method: Compositional matrix adjust.
 Identities = 10/23 (43%), Positives = 15/23 (65%), Gaps = 0/23 (0%)

Query  34   KPIIGVANGFSTITPCNMGLNKL  56
            KP I  A+GF+ +  C +G+N L
Sbjct  301  KPAILAASGFTVVWICYLGVNLL  323



Lambda      K        H        a         alpha
   0.315    0.135    0.377    0.792     4.96

Gapped
Lambda      K        H        a         alpha    sigma
   0.267   0.0410    0.140     1.90     42.6     43.6

Effective search space used: 390292080


  Database: /bio/db/fasta/genes/T00004.pep
    Posted date:  Dec 7, 2016  9:25 AM
  Number of letters in database: 1,136,958
  Number of sequences in database:  3,564
```

```
Matrix: BLOSUM62
Gap Penalties: Existence: 11, Extension: 1
Neighboring words threshold: 11
Window for multiple hits: 40
```
