## Supplemental material 2 for "Dynamic allocation of carbon storage and nutrient-dependent exudation in a revised genome-scale model of *Prochlorococcus*"

### Independent Section

Contains tests that are independent of the class of modeled organism, a model's complexity or types of identifiers that are used to describe its components. Parameterization or initialization of the network is not required. See readme for more details.

#### Consistency

| | |
|---|---|
| Stoichiometric Consistency | 34.4% (X3) |
| Mass Balance | 76.2% |
| Charge Balance | 100.0% |
| Metabolite Connectivity | 100.0% |
| Unbounded Flux In Default Medium | 85.8% |
| Sub Total | 66% (X3) |

#### Annotation - Metabolites

| | |
|---|---|
| Presence of Metabolite Annotation | 0.0% |
| Metabolite Annotations Per Database | Info |
| pubchem.compound | 0.0% |
| kegg.compound | 0.0% |
| seed.compound | 0.0% |
| inchikey | 0.0% |
| inchi | 0.0% |
| chebi | 0.0% |
| hmdb | 0.0% |
| reactome | 0.0% |
| metanetx.chemical | 0.0% |
| bigg.metabolite | 0.0% |
| biocyc | 0.0% |
| Metabolite Annotation Conformity Per Database | Info |
| pubchem.compound | 0.0% |
| kegg.compound | 0.0% |
| seed.compound | 0.0% |
| inchikey | 0.0% |
| inchi | 0.0% |
| chebi | 0.0% |
| hmdb | 0.0% |
| reactome | 0.0% |
| metanetx.chemical | 0.0% |
| bigg.metabolite | 0.0% |
| biocyc | 0.0% |
| Uniform Metabolite Identifier Namespace | 0.0% |
| Sub Total | 0% |

#### Annotation - Reactions

| | |
|---|---|
| Presence of Reaction Annotation | 0.0% |
| Reaction Annotations Per Database | Info |
| rhea | 0.0% |

### Specific Section

Covers general statistics and specific aspects of a metabolic network that are not universally applicable. See readme for more details.

#### SBML

| | |
|---|---|
| SBML Level and Version | SBML Level 3 Version 1 |
| FBC enabled | true |

#### Basic Information

| | |
|---|---|
| Model Identifier | COBRA Model |
| Total Metabolites | 802 |
| Total Reactions | 994 |
| Total Genes | 595 |
| Total Compartments | 5 |
| Metabolic Coverage | 1.67 |

#### Metabolite Information

| | |
|---|---|
| Unique Metabolites | 802 |
| Duplicate Metabolites in Identical Compartments | 0 |
| Metabolites without Charge | 0 |
| Metabolites without Formula | 0 |
| Medium Components | 24 |

#### Reaction Information

| | |
|---|---|
| Purely Metabolic Reactions | 790 |
| Purely Metabolic Reactions with Constraints | 5 |
| Transport Reactions | 101 |
| Transport Reactions with Constraints | 2 |
| Thermodynamic Reversibility of Purely Metabolic Reactions | 1.00 |
| Reactions With Partially Identical Annotations | 0.00 |
| Duplicate Reactions | 0.00 |
| Reactions With Identical Genes | 0.44 |

#### Gene-Protein-Reaction (GPR) Associations

| | |
|---|---|
| Reactions without GPR | 164 |
| Fraction of Transport Reactions without GPR | 0.76 |
| Enzyme Complexes | 190 |

#### Biomass

| | |
|---|---|
| Biomass Reactions Identified | 2 |
| Biomass Consistency | Info |
| BIOMASS | 0.00 |
| BiomassTRANS | 0.00 |
| Biomass Production In Default Medium | Info |
| BIOMASS | 0.10 |

| | | |
|---|---|---|
| metanetx.reaction | 0.0% | |
| bigg.reaction | 0.0% | |
| reactome | 0.0% | |
| ec-code | 0.0% | |
| brenda | 0.0% | |
| biocyc | 0.0% | |
| Reaction Annotation Conformity Per Database | | Info |
| rhea | 0.0% | |
| kegg.reaction | 0.0% | |
| seed.reaction | 0.0% | |
| metanetx.reaction | 0.0% | |
| bigg.reaction | 0.0% | |
| reactome | 0.0% | |
| ec-code | 0.0% | |
| brenda | 0.0% | |
| biocyc | 0.0% | |
| Uniform Reaction Identifier Namespace | 100.0% | |

Sub Total 25%

### Annotation - Genes

| | | |
|---|---|---|
| Presence of Gene Annotation | 0.0% | |
| Gene Annotations Per Database | | Info |
| refseq | 0.0% | |
| uniprot | 0.0% | |
| ecogene | 0.0% | |
| kegg.genes | 0.0% | |
| ncbigi | 0.0% | |
| ncbigene | 0.0% | |
| ncbiprotein | 0.0% | |
| ccds | 0.0% | |
| hprd | 0.0% | |
| asap | 0.0% | |
| Gene Annotation Conformity Per Database | | Info |
| refseq | 0.0% | |
| uniprot | 0.0% | |
| ecogene | 0.0% | |
| kegg.genes | 0.0% | |
| ncbigi | 0.0% | |
| ncbigene | 0.0% | |
| ncbiprotein | 0.0% | |
| ccds | 0.0% | |
| hprd | 0.0% | |
| asap | 0.0% | |

Sub Total 0%

### Annotation - SBO Terms

| | | |
|---|---|---|
| BIOMASS | false | |
| BiomassTRANS | false | |
| Biomass Production In Complete Medium | | Info |
| BIOMASS | 54.20 | |
| BiomassTRANS | 54.20 | |
| Blocked Biomass Precursors In Default Medium | | Info |
| BIOMASS | 0 | |
| BiomassTRANS | 0 | |
| Blocked Biomass Precursors In Complete Medium | | Info |
| BIOMASS | 0 | |
| BiomassTRANS | 0 | |
| Ratio of Direct Metabolites in Biomass Reaction | | Info |
| BIOMASS | 0.00 | |
| BiomassTRANS | 0.00 | |
| Number of Missing Essential Biomass Precursors | | Info |
| BIOMASS | 37 | |
| BiomassTRANS | 37 | |

### Energy Metabolism

| | | |
|---|---|---|
| Non-Growth Associated Maintenance Reaction | Errored | |
| Growth-associated Maintenance in Biomass Reaction | | Info |
| BIOMASS | false | |
| BiomassTRANS | false | |
| Number of Reversible Oxygen-Containing Reactions | 8 | |
| Erroneous Energy-generating Cycles | | Info |
| MNXM3 | Skipped | |
| MNXM63 | Skipped | |
| MNXM51 | Skipped | |
| MNXM121 | Skipped | |
| MNXM423 | Skipped | |
| MNXM6 | Skipped | |
| MNXM10 | Skipped | |
| MNXM38 | Skipped | |
| MNXM208 | Skipped | |
| MNXM191 | Skipped | |
| MNXM223 | Skipped | |
| MNXM7517 | Skipped | |
| MNXM12233 | Skipped | |
| MNXM558 | Skipped | |
| MNXM21 | Skipped | |
| MNXM89557 | Skipped | |

### Network Topology

| | |
|---|---|
| Universally Blocked Reactions | 79 |
| Orphan Metabolites | 9 |

| | |
|---|---|
| Reaction General SBO Presence | 0.0% |
| Metabolic Reaction SBO:0000176 Presence | 0.0% |
| Transport Reaction SBO:0000185 Presence | 0.0% |
| Exchange Reaction SBO:0000627 Presence | 0.0% |
| Demand Reaction SBO:0000628 Presence | Skipped |
| Sink Reactions SBO:0000632 Presence | Skipped |
| Gene General SBO Presence | 0.0% |
| Gene SBO:0000243 Presence | 0.0% |
| Biomass Reactions SBO:0000629 Presence | 0.0% |
| Sub Total | 0% |

X2

| | |
|---|---|
| Total Score | 28% |

Total Score

# 28%

Score per Category

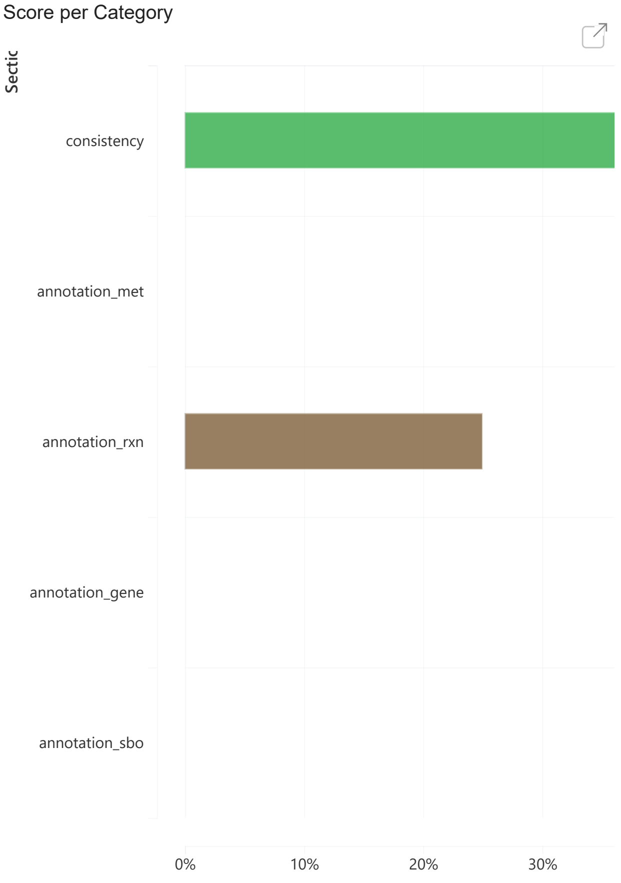


| | |
|---|---|
| Metabolite Production In Complete Medium | 101 |
| Metabolite Consumption In Complete Medium | 103 |

### Matrix Conditioning

| | |
|---|---|
| Ratio Min/Max Non-Zero Coefficients | 0.00 |
| Independent Conservation Relations | 35 |
| Rank | 767 |
| Degrees Of Freedom | 227 |

### Experimental Data Comparison

| | |
|---|---|
| Growth Prediction | Skipped |
| Gene Essentiality Prediction | Skipped |

### Misc. Tests

### Environment

| | |
|---|---|
| Python Version | 3.7.3 |
| Platform | Windows |
| Memote Version | 0.9.13 |
